# Pan-tissue Transcriptome Analysis Reveals Sex-dimorphic Human Aging

**DOI:** 10.1101/2023.05.26.542373

**Authors:** Siqi Wang, Danyue Dong, Xin Li, Zefeng Wang

## Abstract

Complex diseases often exhibit sex-dimorphism in morbidity and prognosis, many of which are age-related. However, the underlying mechanisms of sex-dimorphic aging remain foggy, with limited studies across multiple tissues. We systematically analyzed ∼17,000 transcriptomes from 35 human tissues to quantitatively evaluate the individual and combined contributions of sex and age to transcriptomic variations. We discovered extensive sex-dimorphisms during aging with distinct patterns of change in gene expression and alternative splicing (AS). Intriguingly, the male-biased age-associated AS events have a stronger association with Alzheimer’s disease, and the female-biased events are often regulated by several sex-biased splicing factors that may be controlled by estrogen receptors. Breakpoint analysis showed that sex-dimorphic aging rates are significantly associated with decline of sex hormones, with males having a larger and earlier transcriptome change. Collectively, this study uncovered an essential role of sex during aging at the molecular and multi-tissue levels, providing insight into sex-dimorphic regulatory patterns.

## Introduction

Human diseases often exhibit differences between females and males, including sex-differential morbidity, prognosis, and mortality ^1–3^. The sex-dimorphism has been widely reported in neurological disorders ^4,5^, cardiovascular diseases ^6^, immunological defects ^7^, and cancers ^8^. Accordingly, the life expectancy shows substantial variability between females and males. The evidence on sex-differential mortality was recently reported during the COVID-19 pandemic, mainly affected by different immune responses between females and males ^9,10^. The known molecular mechanisms mostly revolved around sex-differential genetic variants, epigenetics, transcriptomes, and sex-differential responses to environmental exposures ^2^. It is worth noting that many diseases with sex-dimorphism are also age-related, especially in neurodegenerative disorders, cardiovascular diseases, and cancers ^11^. Such sex-differential susceptibility contributes substantially to different life expectancies between females and males, and the related underlying mechanisms are thought to be hormone-driven, mitochondria-related, and sex-chromosome-linked ^11–15^. Previously, sex-differential aging signatures, including chronological trends and gene networks, have been studied in the whole blood and brain regions of healthy donors ^16^, revealing the sex differences of disease vulnerabilities during aging ^17,18^. Sex differences regarding the composition and inflammatory signaling of immunocytes in blood were also analyzed in single-cell resolution, which suggested sex-differential aging in the immune response ^19^. Moreover, at proteogenomic level, sex-biased genes play key roles in several important cellular processes during cardiac aging, such as mitochondrial metabolism, RNA splicing, and translation, implying sex dimorphism in cardiac diseases ^20^. However, a systematic analysis across multiple tissues on the sex-differential aging and underlying molecular mechanisms is currently lacking.

The Genotype-Tissue Expression (GTEx) project contains a large set of high-throughput sequencing data from postmortem donors across 54 human tissues ^21,22^. Previous studies have uncovered many sex-biased genes or expression quantitative trait loci (eQTLs)^22,23^ across different tissues that could affect several critical cellular signal pathways related to human diseases. Since the GTEx donors have a wide age range of 20-70 years, it is possible to identify genes and molecular pathways significantly changed during aging process ^24,25^. However, the relationship between sex-differential regulation and aging has not been thoroughly studied on a transcriptome or pan-tissue level scale. Therefore, the GTEx data could also provide an integrated resource for a comprehensive study of this topic.

In addition to transcriptional regulation, most human genes undergo alternative splicing (AS) to produce different isoforms with distinct activities ^26^. AS is tightly regulated in various tissues or developmental stages by multiple *cis*-acting elements and *trans*-acting factors ^27,28^. Dysregulation of AS is closely related to many age-related diseases, such as neurodegenerative diseases ^29^ and cancers ^30^. Since most introns are co-transcriptionally spliced, AS is also affected by various factors that regulate transcription, such as promoter activity, Pol II elongation, and chromatin modification/remodelling ^28^. Recent studies reported that AS regulation is closely associated with sex ^31^ and age ^32–34^; thus, it is intriguing to further explore the AS regulation during aging stages in different sexes.

In this study, we initiated an integrated analysis of pan-tissue transcriptome data in GTEx, and systematically determined how sex and age contributed to the global variations of gene expression (GE) and AS (Fig. 1). We further focused on the sex-biased AS changes during aging, and found that these AS events are significantly associated with neurodegenerative diseases in a sex-biased fashion. We further examined the chronological changes of age-associated genes, and found a sex-dimorphic transcriptome aging, with males showing an earlier onset of aging and a faster aging rate in most tissues. Collectively, this study clarified the sex and age effects on the transcriptomic changes, revealing the sex-dimorphic aging at a multi-tissue level and their potential mechanisms.

**Fig. 1.**
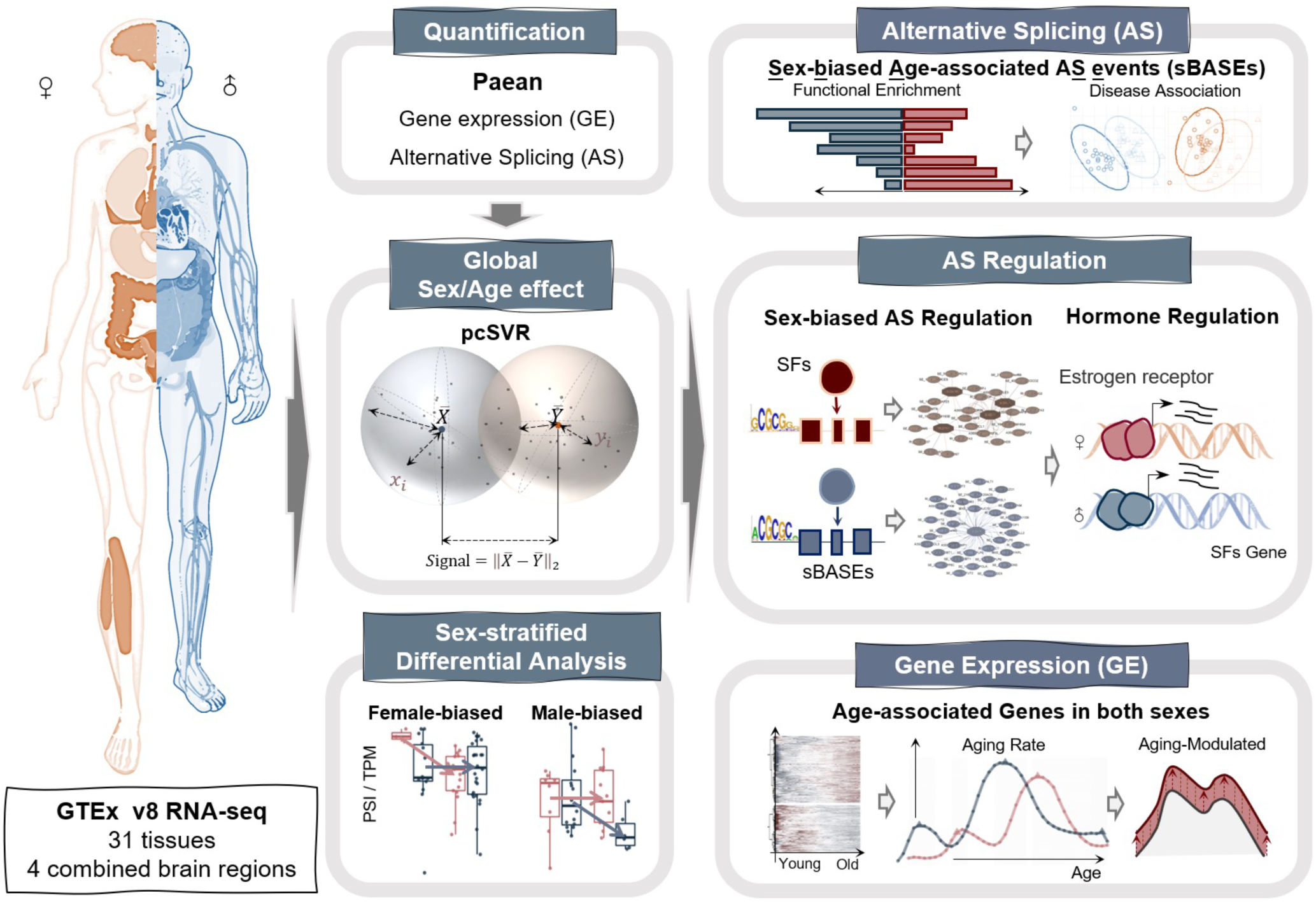
Schematic Overview of our study.

## Results

### Sex and age are critical drivers to the global transcriptome variation

To systematically study how sex and age affect transcriptome complexity, we conducted a thorough analysis of RNA-seq data from GTEx project ^21^ to quantify the GE and AS patterns of all genes from 54 tissues (Fig. 1). Due to data sparseness, the brain tissues were recombined into four functional regions (fig. S1, table S1), including hormone- or emotion-related region, movement-related region, memory-related region, and decision-related region (See Methods). We performed the principal component analysis (PCA) on GE and AS data, and observed the sex and age differences in several tissues (fig. S2). Based on a previous study ^35^, we designed a method called <u>p</u>rincipal <u>c</u>omponent-based <u>s</u>ignal-to-<u>v</u>ariation <u>r</u>atio (pcSVR) to measure the adjusted Euclidean distance in higher dimensions. The pcSVR quantified the distance between different sex or age groups divided by the data dispersion within each group (intrinsic differences between groups will result in a pcSVR value significantly larger than 1), serving as a reliable measurement for the sex or age effects on transcriptomic variations (fig. S3A, see Materials and Methods, Equations 1-3). Compared with the direct identification of differentially expressed genes or AS events, pcSVR provides a global measurement by considering variations from all genes and AS events between different groups.

We grouped the samples into male *vs.* female and young *vs.* old, and calculated the sex-pcSVR and age-pcSVR in multiple tissues (Fig. 2, A and B). Due to a large variation of menopause ages among different individuals and a continuous decline of sexual steroid levels between ages 40 and 60 ^36,37^, we grouped the samples into young (age<40) *vs.* old (age>60) with an age gap instead of a specific age cutoff to reduce the data noise. We found that the breast tissue showed the largest sex-pcSVR, while the adipose and cardiovascular tissues showed larger age-pcSVR (i.e., differences between age groups). The tissues with significant sex differences were similar to those reported in a previous study of sex-differential GE using transcriptomic signal-to-noise ratio ^35^, including breast, thyroid, skin, and adipose. Using the permutation test for pcSVRs, we found that age showed substantially larger effects than sex to human transcriptome in most tissues as judged by both GE and AS (Fig. 2, A and B, table S2). Moreover, AS was significantly affected by both sex and age across most tissues, while GE was affected by sex in a much smaller number of tissues as compared to AS profiles (i.e., comparing the orange triangles in Fig. 2A *vs.* 2B). For example, the coronary artery and adrenal gland are significantly affected by sex and age in their AS profile, but their GE profiles are not affected by sex or age (Fig. 2, A and B). Similar results were found even when removing the genes and AS events encoded by sex-chromosomes (fig. S3B and C), selecting a range of PC cutoffs to capture different global variance (fig. S3, D and E), or using other approaches for p-value estimation (e.g., boot-strapping with replacement, see Materials and Methods, fig. S3, F and G).

**Fig. 2.**
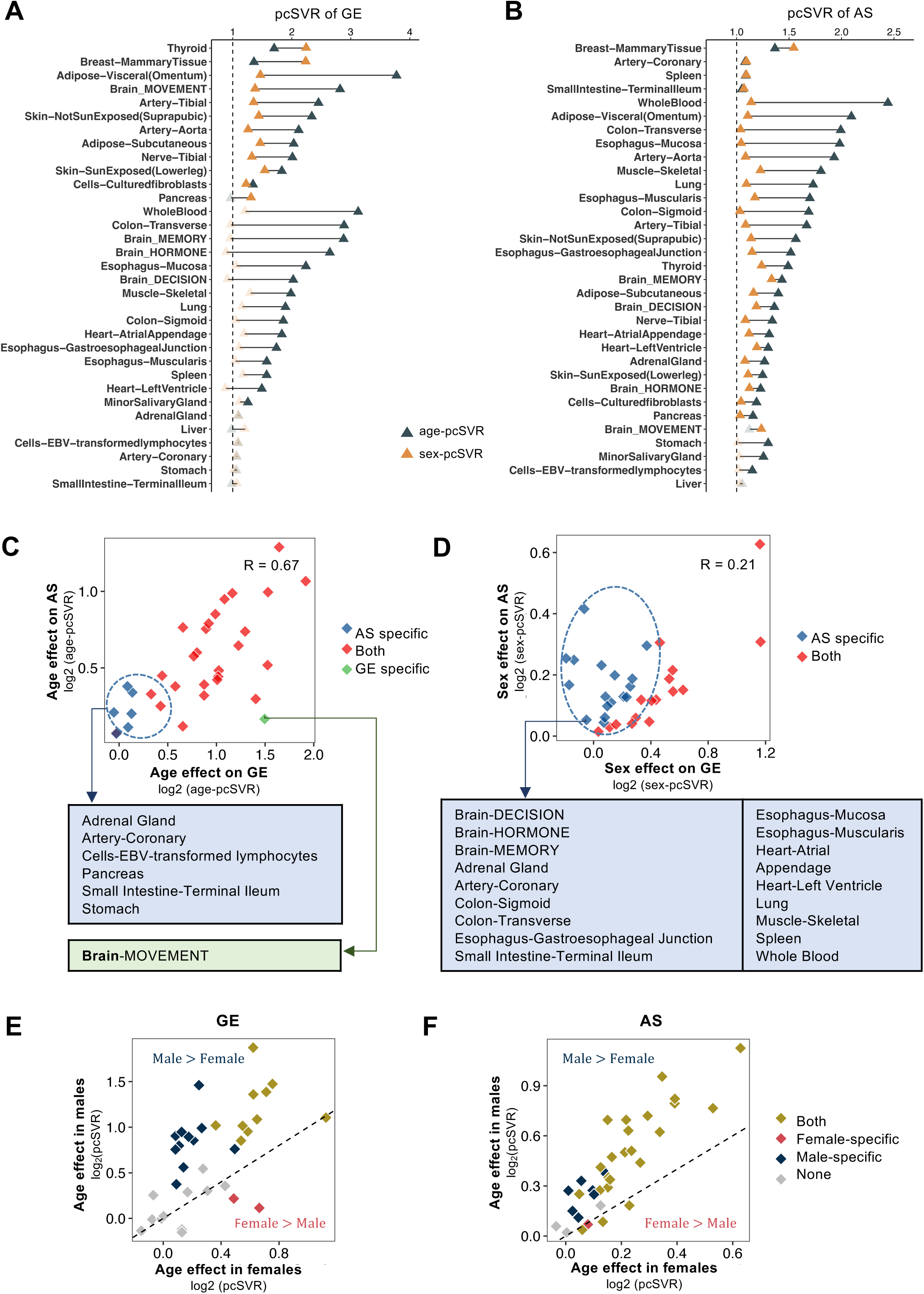
Individual sex or age effects and combined age-by-sex effects on global transcriptomic variation.

We further compared the age and sex effects calculated by GE *vs.* AS (table S2), and found a significant correlation between the age-pcSVR of GE and AS in most tissues (R=0.67, p<0.05, Fig. 2C), suggesting the age contribution to transcriptome variation is largely consistent regardless of the input data types (i.e., GE or AS). However, the sex effects on GE and AS showed a much weaker correlation (R=0.21, Fig. 2D), consistent with our earlier results (Fig. 2, A and B). We also found that age was a more vital driver of AS variation than GE in specific tissues like coronary artery and stomach, whereas sex contributed more to AS variations in whole blood, skeletal muscle, and most brain regions.

To further examine the sex-differential age effect, we compared the age-pcSVR between females and males (Fig. 2, E and F; GE and AS, respectively). Consistent with earlier findings (Fig. 2C), we found significant age effects in both sexes across most tissues. Moreover, the age effects in both GE and AS were apparently larger in males than in females (Fig. 2, E and F). These results suggested that age and sex contributed significantly to transcriptome variation in various tissues with certain degrees of heterogeneity for different data types.

### Differential GE and AS analyses reveal general interactions of age-by-sex

We next identified specific genes or AS events responsible for sex or age effect on transcriptomic variations *via* a linear regression model containing an age-by-sex coefficient term (see Materials and Methods, Equations 4 and 5). In each tissue, we identified the sex- or age-differential genes and AS events (fig. S4, A and B). Consistent with previous reports ^35,38,39^, the breast tissue contains the largest amount of sex-differentially expressed genes, while the age-biased genes were widely discovered in brain regions and artery-related tissues. Interestingly, in most tissues, there were no significant overlaps between age-differential genes and AS events (fig. S4A, hypergeometric test p-value < 0.01). In contrast, we found significant overlaps between sex-differential genes and AS events in most tissues (fig. S4B).

As a control, we constructed a separate regression model to evaluate the contribution of sampling biases, including sample sizes and the total number of genes or AS events detected in each tissue (Equation 6). We found that the sample size did not significantly affect the number of identified sex-differential or age-differential genes/AS events (fig. S4, C and D), suggesting our analyses are not affected by sampling biases. Furthermore, using the coefficient of the age-by-sex term in this model allowed us to identify thousands of genes and AS events affected by the functional interactions between sex and age (i.e., the sex effects on GE/AS depend on the age, *or vice versa*) (fig. S4E), suggesting extensive sex-dimorphic changes of the transcriptome during aging.

A recent study reported the environmental effect on human genetic background during aging ^40^, which should be considered when analyzing large datasets from different populations. In the linear regression model, we included the surrogate variables (SV) as confounders to control for the effect of genetic backgrounds (Equations 4 and 5). Using the differential GE analysis in whole blood as an example (other tissues have the same trend), we evaluated the correlation between SVs with donors’ ethnicity using Point Biserial Correlation and found that many SVs showed significant correlations with the donors’ ethnical background (fig. S5A), suggesting that these SVs could effectively reflect the variables of genetic background. Additionally, we calculated the correlation between SVs with the top 5 principal components as judged by whole genome sequencing (WGS) data (fig. S5B), and found roughly the same set of SVs also showed a significant correlation with PC1.

Moreover, recent studies demonstrated the powerful impacts of post-mortem interval (PMI) and time of death (TOD), which included the death seasons and day times on gene expression ^41,42^. Thus, we carefully evaluated whether both factors are controlled as potential confounders in our linear model. Our results showed that PMI and TOD significantly correlate with covariates in most tissues (fig. S5C-E), suggesting that their effects could be effectively regressed in our model. Together, the SVs in the linear regression models could effectively control the cofounders of human genetic regulation. In addition, we did the differential analysis that incorporated PMI as a covariate in the regression models and re-evaluated the age- and sex-related transcriptomic changes. Using Whole Blood gene expression as an example, our revised analysis shows that the inclusion of PMI in the covariates has minimal impact on the significance levels and effects of sex and age (i.e., p-values and coefficients, respectively), indicating that our findings are robust using confounding factors (fig. S5F-G).

To further evaluate the robustness of the differential analyses, we performed detailed assessments of the linear regression models and the selection of GE/AS cutoffs using the decision-related brain region as an example. We calculated the fraction of original differential genes/AS events identified in each permutation iteration, and the false discovery rate (FDR) for each gene/AS event using the permutation analysis in the linear regression model (see Materials and Methods). Briefly, we generated the permuted data 1,000 times by randomly shuffling the group labels (sex and age) with the same sample sizes as the original labels. The distribution of the p-values of sex/age-differential genes/AS events identified based on the original and each shuffled phenotype data were evaluated (fig. S6, A and D). We first observed low fractions of sex/age-differential genes/AS events in most iterations under our cutoffs (fig. S6, B and E). We found that FDRs of most sex/age-differential genes/AS events were not more than 5% using the linear regression model (fig. S6, C and F), which is much lower than the FDR of DESeq2 and edgeR ^43^. As a result, this approach could justifiably control the false discovery for differential analysis. Furthermore, the TPMs of most sex- or age-differential genes were higher than 1, and the junction read counts (JC) of most differential AS events were higher than 10 (fig. S7, A and B), suggesting that our cutoffs (TPM > 1 or JC > 10) preserved most of the differential GE and AS events. In addition, we found a low overlap between the genes with age-differential GE and AS under most combinations of TPM and JC cutoffs in the decision-related brain region (fig. S7C), which is consistent with our earlier findings (fig. S4A). Collectively, these controls confirmed the reliability of our differential analyses.

### AS regulation shows a stronger sex-dimorphism during aging than GE across all tissues

We further studied the transcriptomic changes during aging process in different sexes by identifying age-associated genes or AS events separately in females and males (table S3). We found that the age-associated genes significantly overlapped between females and males in most tissues (p-values < 0.01 by hypergeometric tests, Fig. 3A), whereas the overlaps of the age-associated AS events were much smaller (Fig. 3B). Consistently, in most tissues, the age effect on GE was more correlated between females and males (Spearman’s correlation between β_F_ and β_M_, Equation 7) than the age effect on AS (Fig. 3C), regardless of different GE/AS filters used in such analysis (fig. S7D). Taken together, these observations suggested that the age effect on GE is more consistent in both sexes, while age-associated AS showed a stronger discrepancy between two sexes, implying that splicing regulation had a stronger sex-dimorphism during aging.

**Fig. 3.**
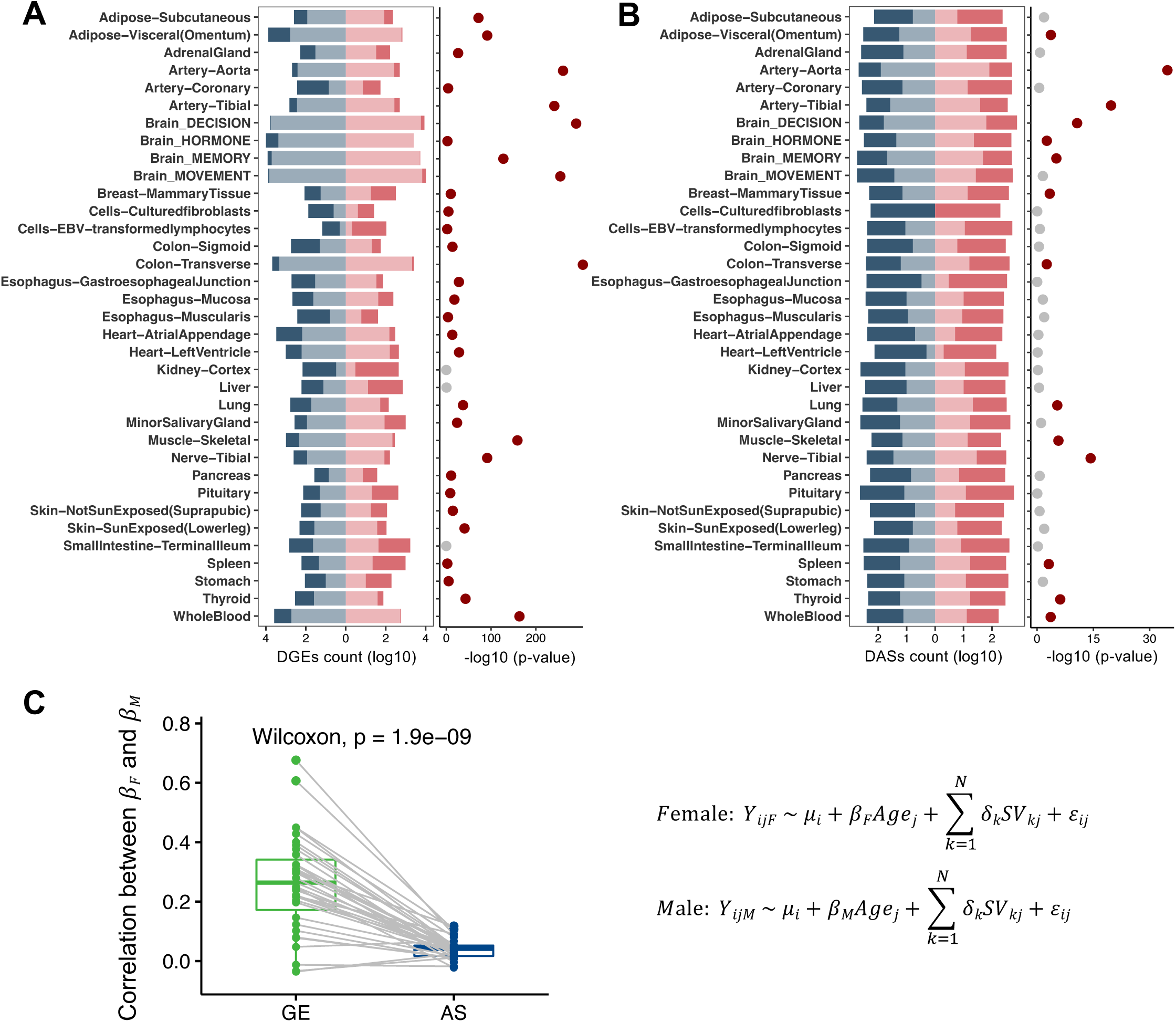
Sex-stratified age-associated genes and AS events across tissues.

### AS changes show a male-biased association with Alzheimer’s disease in human brain

Because most age-associated AS events were sex-specific, we defined the AS events significantly affected by age in only one sex as the <u>s</u>ex-<u>b</u>iased <u>a</u>ge-associated A<u>S</u> <u>e</u>vents (sBASEs), and further examined their functional impacts on human health using Gene Ontology and Disease Ontology analyses. The functional association of sBASEs varied considerably in different tissues and sexes (fig. S8, table S4). Having established patterns across all tissues, we next focused on tissue-specific AS changes in the brain that is highly affected by age-related processes. Since many neurological conditions show sex differences, the sex-biased splicing events in the brain could be helpful to explain differential susceptibility to age-related cognitive decline and neurodegeneration. Notably, the sBASEs in brain regions were significantly enriched in pathways such as amyloid-beta formation and cytoskeleton (Fig. 4A), with the male decision-related brain region being particularly associated with cognitive and psychotic disorders (fig. S9A). Because previous studies suggested that the disease manifestations of Alzheimer’s Disease (AD) showed sex differences in cognitive decline and brain atrophy ^4^, we next focused on sBASEs in brain to explore their contribution to the sex-dimorphic risks of neurodegenerative diseases. This question was addressed by an integrative analysis of the large datasets from other independent AD studies, which is a common strategy (and strength) of computation biology in the big data era.

**Fig. 4.**
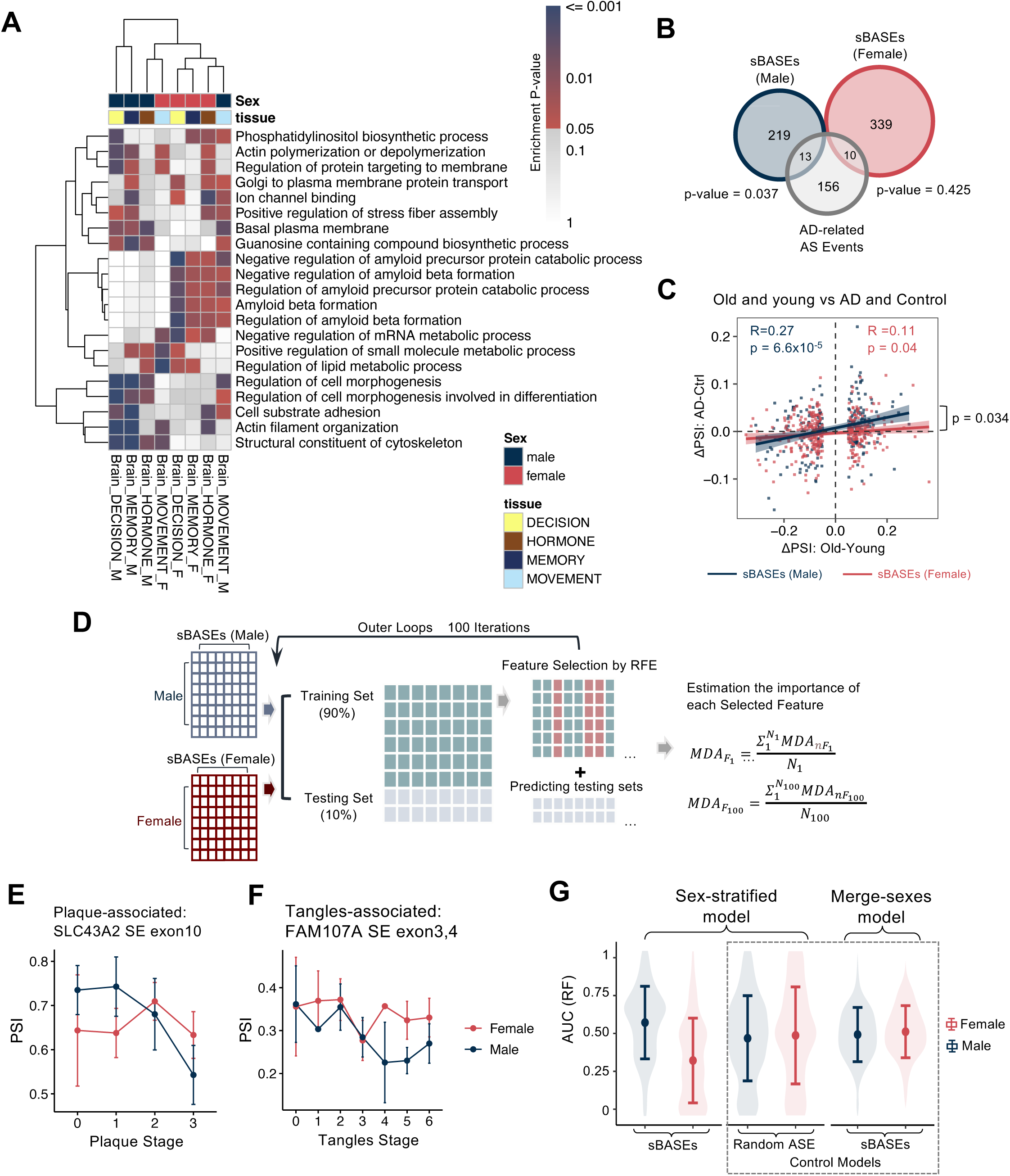
Male-biased associations between the AS changes during aging and neurodegenerative disease.

We analyzed the RNA-seq dataset of the brain prefrontal cortex with a dataset of 91 AD patients and healthy controls manually curated to match the sex and disease ratio (Female:Male≈1:1; AD:Control≈1:1, see Materials and Methods). Surprisingly, although the number of sBASEs in males was smaller compared with that in females, we found that the AD-related AS events significantly overlapped with the sBASEs in males (p = 0.037) but not in females (p = 0.425) (Fig. 4B). In addition, the magnitude of AD-associated AS changes across the male-specific sBASEs (i.e., *PSI_AD_* – *PSI_ctrl_*) was significantly correlated with the age-associated AS changes in males (i.e., *PSI_old_* – *PSI_young_*) (Pearson’s product-moment correlation, R=0.27, P=6.6×10^−5^), and such correlation was significantly higher than that in a separate analysis in females using the female-specific sBASEs (R=0.11, P=0.04; Fig. 4C; see Materials and Methods). In contrast, we found a significant overlap between the AD-related genes and the sex-biased age-associated GE changes in females (p = 0.031) but not in males (p = 0.60) (fig. S9B), supporting a recent finding that females are more vulnerable to AD during aging due to the post-menopause activation of C/EBPβ–AEP/δ-secretase pathway by sex hormone changes ^44^.

In addition, we performed similar analyses using an independent dataset from ROSMAP containing 271 filtered RNA-seq samples from dorsolateral prefrontal cortex (DLPFC) with matched age distribution between AD and control samples. As expected, we found that the AD-associated AS changes of sBASEs in males showed a significantly stronger positive correlation with the age-associated changes (R=0.509, p < 2.2×10^−16^) than such correlation across the sBASEs in females (R=0.371, p < 2.2×10^-16^) (fig. S9C), which is largely consistent with the earlier finding using a smaller dataset (Fig. 4C). This male-biased association indicated that the AS changes during aging could contribute more to the AD in males than females, suggesting additional molecular complexity in the sex effect of AD.

To further examine the contribution of individual sBASE to AD risk, we constructed a sex-stratified random forest classifier using a subset of sBASEs that were also detected in ROSMAP data for AD prediction. For each sex, we randomly divided the datasets of AD patients and controls into separate training and test sets, and repeated the prediction 100 times (Fig. 4D). For each selected sBASE, we also evaluated its averaged feature importance (i.e., mean decrease accuracy) across 100 iterations and the association with the classic AD neuropathology (such as tangles and plaques), resulting in the identification of a series of key sBASEs significantly associated with AD in a sex-specific manner. For example, the skipping of exon 10 in SLC43A2 negatively correlated with the neuritic plaques stage in males but not in females (Fig. 4E). Meanwhile, skipping of exons 3 and 4 in FAM107A, a gene related to synaptic and cognitive functions, showed a male-specific negative association with neurofibrillary tangle stages (Fig. 4F). In addition, the prediction models based on sBASEs performed significantly better in males than females (Fig. 4G and fig. S9D, left panel), suggesting that age-associated AS events could serve as a better predictor for AD risk in male patients. As negative controls, we performed the same analysis to predict male and female patients separately using randomly selected AS events (sex-stratified model; Fig. 4G, middle panel), and predict all patients with male- or female-specific sBASEs (merge-sexes model; Fig. 4G, right panel). We found that both control models showed no differences in males and females, indicating the crucial roles of sBASEs in AD during male aging. Collectively, our analyses of human brains revealed a surprising sex-biased association between RNA splicing and AD, suggesting that regulation of AS may play a critical role in the sex-dimorphisms of age-related diseases such as AD.

### sBASEs are regulated by splicing factors with sex-dimorphic expression during aging

AS is generally regulated by splicing regulatory *cis*-elements that specifically recruit various *trans*-acting splicing factors (SF) to promote or inhibit splicing of adjacent exons ^27,45^. Dysfunction of the SFs has been reported as a hallmark of aging, presumably by regulating the genes in cellular senescence during aging process ^33^. Consistently, our analyses identified a large number of age-associated SFs in human brain (table S5). To further examine the sex-dimorphic regulation through age-associated SFs, we constructed an SF-RNA regulatory network between age-associated SFs and sBASEs in each sex. This process integrated the correlations between the SFs levels and PSI of sBASEs, the significant changes of sBASEs upon SFs knockdown, and the existence of SFs binding motifs around sBASEs (Fig. 5A, left panel). We required the regulatory connections between the SFs and AS events that pass all these three criteria. Such regulatory networks can be generated in different tissues, and we focused on the brain due to its association with age-related diseases such as cognitive disorders and synaptic diseases.

**Fig. 5.**
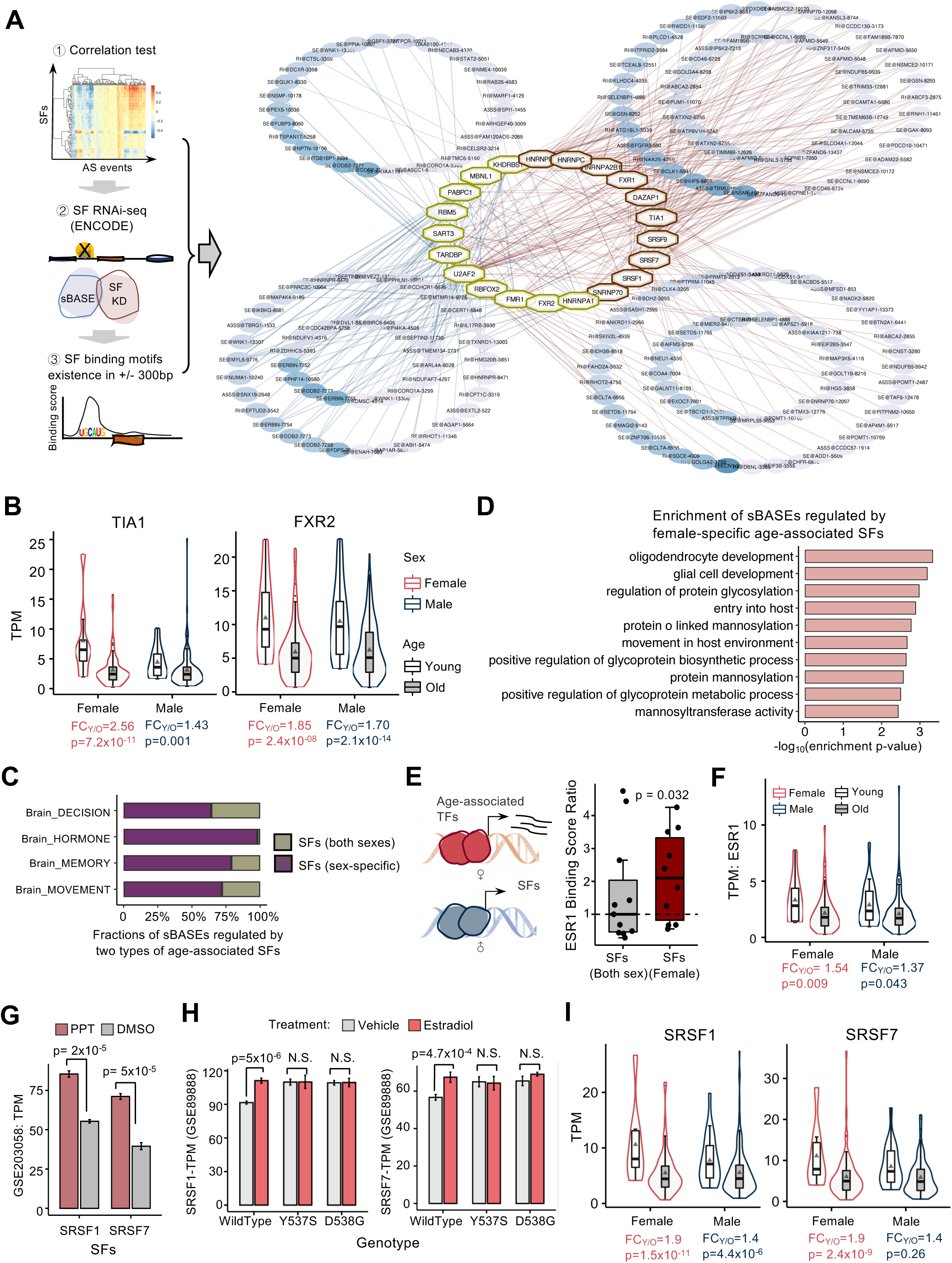
Sex-dimorphic AS regulation during aging in decision-related brain region.

We identified several key age-associated SFs that regulate multiple sBASEs in decision-related brain region (Fig. 5A, right panel). Interestingly, several SFs showed significant age association only in females (e.g., TIA1 in Fig. 5B, left panel), while others were associated with age in both sexes (e.g., FXR2 in Fig. 5B, right panel). This finding is consistent with our earlier observation that more sex-specific age-associated AS events were identified in decision-related brain region of females than males (Fig. 4B). Most of the sBASEs were regulated by sex-specific age-associated SFs rather than the SFs associated with aging in both sexes (Fig. 5C). Functional enrichment analysis showed that these sBASEs regulated by female-specific age-associated SFs were significantly enriched in glial cells and oligodendrocytes development (Fig. 5D), which participated in the myelin generation of aging brain and neurodegenerative diseases in central nervous system ^46^. These results indicated that the SFs with sex-dimorphic expression during aging, especially the female-specific changes of SFs, were one of the main factors in regulating sBASEs with important biological functions.

Furthermore, we studied the potential mechanism that controls the female-specific changes of SFs during aging in decision-related brain region. It was previously reported that the female sex hormone estrogen consistently decreases during lifespan ^37,47^; thus we speculated that estrogen may play roles in controlling transcription of SFs during aging through activation of nuclear receptors. To this end, we analyzed the public ChIP-Atlas data to examine the binding of canonical estrogen receptor ESR1 to the promoter region. We found that the ESR1 binding ratios on the female-specific age-associated SFs were significantly higher than the median value of the age-associated SFs common in both sexes (Fig. 5E; p = 0.032), suggesting that ESR1 was more likely to bind the promoters of the female-specific age-associated SFs. In addition, the age-associated decline of ESR1 was more substantial in females than males (Fig. 5F), and some of the female-specific age-associated SFs (e.g., SRSF1 and SRSF7) could be stimulated by propyl pyrazole triol, an agonist of ERα (Fig. 5G) ^48^.

These observations were further confirmed by an independent experiment where the wildtype and two constitutively active mutants (Y537S, and D538G) of ESR1 were engineered into MCF-7 cells using CRISPR-Cas technology ^49^. The resulting cells were treated with estradiol or vehicle controls and subjected to transcriptome profiling by RNA-seq, and the data were reanalyzed using the same pipeline. We found that the basal levels of SRSF1 and SRSF7 were significantly higher in the cells with constitutively active ESR1 (Fig. 5H, comparing samples treated with vehicle controls), and SRSF1 and SRSF7 were further induced by the estradiol treatment only in cells with wildtype ESR1 but not in cells with mutated ESR1 (Fig. 5H, compare the samples treated with estradiol *vs.* control in different cells), indicating that the activation of ESR1 pathway can promote expression of SRSF1 and SRSF7. The similar changes were also found in other splicing factors including HNRNPA2B1 and HNRNPC (fig. S10A). As expected, the expression levels of SRSF1 and SRSF7 showed an obvious decrease during female aging (Fig. 5I).

To further confirm our findings in different tissues, we analyzed the RNA-seq data with ER knockout and estrogen treatments in mouse brain arcuate nucleus (ARCs) ^50^ and found a consistent trend in the expression changes for splicing factors Srsf1, Srsf7, and Hnrnpa2b1 under estradiol (E2) regulation (fig. S10B). Specifically, these splicing factors were activated by estrogen in wild-type mice but exhibited reduced expression in ER knock-out mice. The direct binding of estrogen was also examined through integrating the ESR1 ChIP-seq data ^51^. As expected, a major ESR1 binding peak was found in the promoter regions of Srsf7, which was increased by estrogen treatment (fig. S10C).

Collectively these analyses across multiple datatypes suggested that the female-specific SFs (such as SRSF1 and SRSF7) are more sensitive to the estrogen changes *via* ESR1-mediated pathway, leading to extended sex-biased AS regulations during aging.

### The aging rates of GE show sex-dimorphism during aging across multiple tissues

In the earlier results (Fig. 3A), we found a large proportion of age-associated genes are common in both sexes. Therefore, we asked whether the levels of GE change at the same pace during aging in females and males. The age-associated genes ubiquitously changed in both sexes were analyzed for consistency. We first used the Autoregressive Integrated Moving Average model ^52^ to capture the age-associated genes with significant chronological changes beyond random fluctuations, which are the main contributors to transcriptome dynamics during aging. We further quantified the GE changes between adjacent age windows using breakpoint analysis ^16^ (Fig. 6A, see Materials and Methods). Such analyses were conducted across all tissues in both sexes (fig. S11A), with two tissues shown in Fig. 6B. A higher value on the Y-axis (mean - log P) indicated a larger GE change at the given age, and thus the peaks were regarded as aging “breakpoints” that represent the key time point of transcriptome aging. Compared to females, males showed significantly larger GE changes during aging in most tissues (fig. S11A), except transverse colon, fibroblast cells, and whole blood (fig. S11B). The number of breakpoints also showed sex-dimorphism in several tissues. For example, in sun-exposed skin and colon, males showed two main breakpoints around ages 35 and 50, whereas females only have one breakpoint at age ∼45 (fig. S11A).

**Fig. 6.**
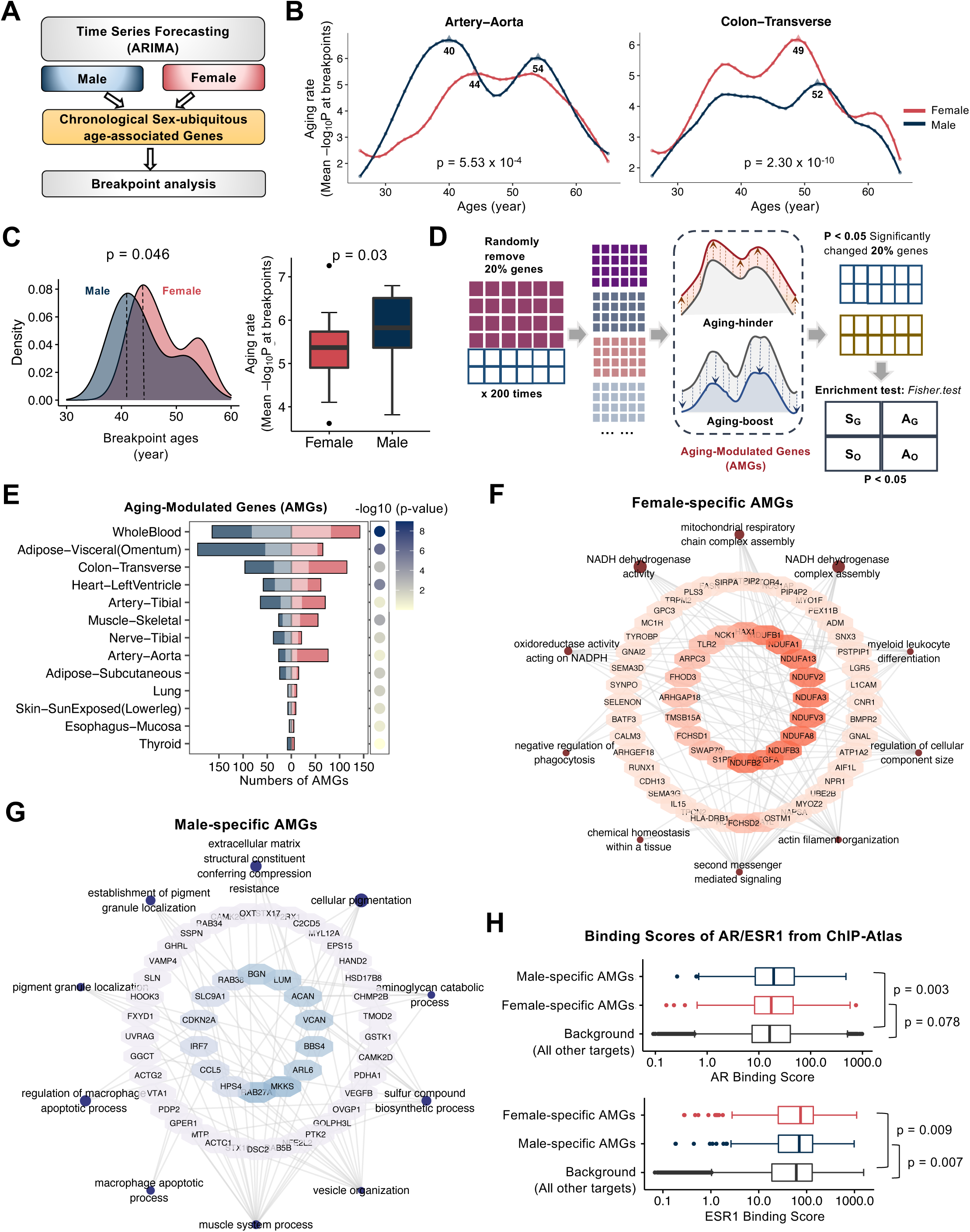
Sex-dimorphic aging rate of GE during aging process.

Moreover, we found that the major aging breakpoints (i.e., breakpoints with the largest GE changes) occurred earlier in males than females across most tissues, and the magnitudes of changes were also more prominent in males (Fig. 6C), suggesting that males may age faster with a larger age-associated GE change. In addition, we conducted breakpoint analysis with all chronological genes in both sexes and found similar results when comparing the breakpoints in males and females (fig. S11C). We also performed a similar analysis using chronological sex-biased age-associated AS events (sBASEs) and all chronological AS events in females and males separately, and again found an earlier and more obvious breakpoint (fig. S11, D and E). In summary, these results uncovered an earlier aging and a higher aging rate in males at molecular levels, again implying that males could be surprisingly more vulnerable to aging.

### Aging contributors in separate sexes show distinct functions

To further examine how the individual gene affected aging rates, we designed a disturbance analysis to identify <u>a</u>ging-modulated <u>g</u>enes (AMGs) (Fig. 6D). Specifically, we randomly removed 20% of chronological genes and repeated the breakpoint analysis with the remaining genes to test if the breakpoints were significantly altered (see Materials and Methods). This process was repeated 200 times, and the genes enriched in the 200 random samplings (judged by Fisher exact test p < 0.05) were defined as AMGs that serve as the key contributors to molecular aging.

We identified a total of 1035 AMGs across different tissues (table S6), around 70% of which (723 in 1035) were sex-specific (Fig. 6E). Functional enrichment analysis suggested that the female-specific AMGs were significantly enriched in cellular respiration and NADH dehydrogenase pathways (Fig. 6F), which is consistent with an earlier report of their roles in human aging and longevity ^53^. Such functional enrichment was not found in the male-specific AMGs (Fig. 6G), implying that female aging may be more sensitive to mitochondrial and oxidative functions. Intriguingly, we also found many male-specific AMGs relevant to myeloid leukocyte differentiation, phagocytosis, and macrophage apoptotic processes, implying a sex-differential role of the immune system during aging.

To determine the potential mechanism of the sex-dimorphic aging rate, we focused on sex hormones that are critical for transcriptional regulation. We found that the androgen receptor (AR) tends to bind the promoters of male-specific AMGs (P=0.007), suggesting potential regulations by androgen during male aging (Fig. 6H, top panel). On the other hand, female-specific AMGs were more likely to be bound and regulated by estrogen receptor ESR1 (Fig. 6H, bottom panel), suggesting the potential regulation by distinct sex hormones.

Furthermore, we examined the detailed expression profile of AMGs from the whole blood that is commonly used for health surveillance. The AMGs common to both sexes can be grouped into three clusters based on age-associated expression profiles. GE of the largest cluster increased at young ages and abruptly decreased in mid-ages in males, but monotonically decreased in females (fig. S12, A and B). These genes were functionally enriched in neutrophil and granulocyte migration and RNA splicing (fig. S12C). Moreover, we also examined sex-specific AMGs and found very distinct expression patterns (fig. S12, D and E). Interestingly, while most of these genes showed decreased expression at old ages, the expression of some male-specific AMGs gradually increased during aging (fig. S12D, dotted box). Taken together, these results suggested that these age-associated genes modulating the transcriptome aging rate showed different functional impacts in females and males.

## Discussion

Previous studies in selected tissues suggested that sex is a key factor in affecting human aging and age-related diseases ^11,12^, which may contribute to different life expectancies between two sexes. However, few pan-tissue investigation on age-by-sex effect was conducted at the transcriptome level, especially in the resolution of AS isoforms. By comprehensively analyzing ∼17,000 samples across 54 tissues, we uncovered individual sex or age effect on global transcriptome variations, as well as the combined age-by-sex effects. Using a sex-stratified differential analysis, we found that AS changes showed a stronger discrepancy between females and males during aging. We further identified the regulatory splicing factors for sBASEs and explored the underlying mechanism of sex-differential AS regulation during aging. Conversely, we found the change of age-associated gene expression was consistent between two sexes, with a faster aging rate in males. Our analysis provided a comprehensive landscape of how sex and age affect human transcriptome, revealing the sex-dimorphic aging at a multi-tissue level.

Multiple mRNA isoforms can be produced from a single gene through AS, increasing the transcriptomic and proteomic complexity in different cells ^54,55^. Therefore, analyses of AS changes during aging of each sex may provide additional information beyond the GE profile. Interestingly, we observed different patterns when analyzing the profiles of GE *vs.* AS. For example, judged by the AS changes, we observed more tissues with a significant sex or age effect on transcriptome and a higher discrepancy between females and males from the combinatorial effects of age-by-sex (Fig. 3), highlighting the contribution of splicing in sex dimorphism during human aging at an isoform-specific resolution.

The sex-dimorphic patterns of AS and GE not only illuminate aging-related transcriptomic changes but also reveal intriguing connections to age-related diseases, such as Alzheimer’s disease (AD). Most studies showed that females have a higher risk of AD due to the post-menopause changes of sex hormones ^44^, however, a higher incidence rate of mild cognitive impairment in males is also reported ^56^. Additionally, based on an early study on a large cohort of 17,127 participants from UK Biobank, age-related cognitive risk factors are more widely discovered in males, suggesting a potential vulnerability of males to cognitive decline with age ^57^. We further validated our findings using Alzheimer’s disease dataset, ROSMAP, where the consistent correlations between aging-related splicing changes and AD-related changes were observed, providing additional evidence for the robustness of our results. At the splicing level, we surprisingly observed a male-biased association between age-associated AS changes and AD (Fig. 4). Such associations could not be detected by analyzing GE, suggesting that analysis at isoform resolution can provide different information into sex-dimorphic aging. We also designed a sex-stratified random forest classifier to predict AD in males and identify several key AS events contributing to such prediction. Due to the limited sample size, large variations were observed between different training/testing iterations during the sex-stratified AD prediction. However, we still observed a higher performance of the classifier in males compared to the control model, which is in concordance with that the AS changes during aging may contribute more to AD in males than females. Our findings suggest that male-biased age-associated AS changes may contribute to the increased vulnerability of males to cognitive decline, providing a complementary perspective to existing research. We expect that future investigations using bigger datasets will further identify the sex differences in AD.

Using a pan-tissues analysis, we discovered that transcriptomic aging happened earlier in males than in females (Fig. 6D), consistent with the reported shorter life expectancy in males ^12^. Furthermore, we identified many sex-specific AMGs with different biological functions, such as the macrophage and mitochondria functions (Fig. 6, F and G). These genes tend to be regulated by the dominant sex hormones in each sex. The investigation of the aging rate provided insights into the intervention of the aging process or age-related diseases, and to some extent made available for identifying therapeutic targets at specific age points. It is worth noting that our analysis of the aging rate in this section does not include the data from brain tissues due to the sample sparsity in selecting parameters.

Our analyses also have some technical limitations, the biggest of which may be caused by the cell type and sample heterogeneity between different donors of GTEx datasets. Since GTEx dataset is the most comprehensive dataset with both transcriptome and whole genome sequencing data, it’s a good public resource for sex, age, and age-by-sex interaction analysis within a healthy context. Compared with the earlier studies that used either microarray or other RNA-seq data to analyze the effect of age or sex on brain transcriptome, our main findings supported the main conclusion of these reports, including the diverse gene profiles across brain regions during aging, as well as more vulnerability in males on a global scale ^17,18^. In addition, GTEx has a very strict selection of healthy donors to reduce biases from different diseases ^58^, which is a unique feature compared to other cohorts that often contain patients of various diseases that may affect the analyses at the molecular level. In particular, the large-scale brain samples provide a unique opportunity to analyze transcriptomic changes in sex-dimorphic aging. However, several technical challenges could limit the generalizability due to dataset-specific biases, including cell type heterogeneity, postmortem artifacts, as well as sequencing biases. For example, GTEx data is bulk RNA-seq, which does not capture cell-type-specific transcriptomic changes. Given the cellular complexity of the brain tissues, the observed differences in gene expression and splicing may be influenced by shifts in cellular composition rather than intrinsic transcriptional regulation. Future studies involving orthogonal datasets and functional validation will be essential to substantiate the biological relevance of the identified sex- and age-associated splicing changes.

The second limitation is the existence of potential confounding factors (like batch effect, genetic background, etc.) that are often associated with big data analyses. However, when such batch effects are entangled with the effect from distinct sex/age groups, it becomes technically challenging to effectively remove such factors without canceling out the contribution of sex/age, which is a common technical difficulty reported also in other analyses ^59^. We take several measurements to reduce the effect of confounding factors. First, we utilized the original TPM/PSI data to aggregate sex/age effects and conducted a permutation analysis to assign an empirical *p*-value to each pcSVR. This approach provided a robust statistical framework for evaluating the significance of sex/age effects while accounting for potential batch effects. Secondly, in the context of the differential analysis, we preserved all possible confounders and identified changes that were more obvious than the effects of these confounders, a strategy that reduces false positive discoveries originating from potential batch effects. Another technical limitation is related to the small number of samples in some tissues, and thus we either omitted them from aging analyses (like pituitary, bladder, and kidney) or combined similar tissues (like brain regions) for detailed analyses.

Moreover, we derived the AS regulatory networks using the multi-dimensional large-scale data from highly proliferative and easily cultivable cancer cell lines, which may not be an ideal model for the post-mitotic cells in human tissues. However, we dissected the RBP-RNA regulatory network by integrating muti-dimensional data obtained through several orthogonal state-of-the-art approaches (Fig. 5A), including the eCLIP-seq, *in vitro* RNA affinity evaluation, RNA-seq with RBP depletion, and chromatin association ^60^. In addition, the RBP-RNA binding relationships seem to be universal, and the distinct splicing outcomes are mainly determined by the tissue-specific expression and activity of different splicing regulatory factors. Therefore, it is possible to use the data in cancer cells to derive a connectivity map of RBP-RNA, and then use the tissue-specific expression of RBPs to refine the regulatory map.

It was reported that there is only a moderate correlation between mRNA and protein abundances for many human genes, mainly because of the variation in translation efficiency of different mRNAs and the additional regulation in protein degradation. Here we constructed the SF-RNA regulation network with the shRNA-seq data that represents the reduction of specific splicing factors in both RNA and protein levels. Such data is more reliable than the correlation of cell-specific expression of RNA and protein, and thus it is feasible to evaluate the consequences of gene silencing by looking at the RNA level. To minimize tissue biases in the RBP-RNA regulation network, we incorporated 3-step processes (see Materials and Methods) including the restrictions of the correlations between RBP expression and AS events within each tissue. Additional large-scale experiments are needed in the future to globally profile the physical and functional interactions between RNA and RBP with a higher accuracy from different tissues.

Mechanistically, we found a prominent role of sex hormones in regulating sex dimorphic aging or age-related diseases. Using a range of RNA-seq and ChIP-seq data, we uncovered a type of female-specific age-associated AS regulation (Fig. 5A), which may be achieved partially through the estrogen-mediated regulation of female-specific splicing factors (Fig. 5, E and I). We further quantified chronological changes of transcriptome dynamics using breakpoint analysis, and observed the sex-dimorphic aging rates across multiple tissues. Our analysis observed the non-linear aging patterns with two breakpoints, which is consistent with recent findings, with some differences in specific age points due to sex differences as well as tissue diversities ^61^. In addition, our aging breakpoints are largely consistent with the changes in circulating sex hormones during aging. The level of testosterone starts to decrease at the age 35–40 in males, while in females, estrogen level starts to decrease at a later age (i.e., ∼50 years old) ^37,47^. These breakpoints could represent key junctures in the aging process that align with the non-linear patterns of aging and disease progression. Furthermore, we identified a series of female-specific and male-specific aging-modulated genes (AMGs), whose functions are consistent with the reported roles of sex hormones. For example, estrogen was reported to participate in the antioxidant system *via* NADPH ^37,47^, which is functionally enriched in female-specific AMGs (Fig. 6F). Conversely, androgen was reported to affect the inflammation in immune response ^37,47^, which is enriched in the male-specific AMGs (Fig. 6G). Consistently, we found that the sex-specific AMGs were more likely to be regulated by androgen receptor in males or estrogen receptor in females (Fig.6H). However, the regulation of sex-biased aging is probably more complicated. Other factors, including mitochondrial functions, genes encoded by sex chromosomes, non-coding RNAs, and transposable elements, may also contribute to the sex-dimorphism during aging ^12,14,62,63^. In addition, it is possible that the genetic variants, including expression QTL (eQTLs) and splicing QTLs (sQTL), can affect GE or AS in a sex-biased or age-associated fashion ^22,40,64^. Future studies of these factors will improve our understanding of the intricate regulation of transcriptome dynamics, providing us a broader horizon at a multi-omics scale.

Various biological aging clocks had previously been constructed based on different molecular markers, including DNA methylome ^65–67^, transcriptome ^68^, circulating immune proteins ^69^, human gut microbiome ^70^, and multi-omics features ^71^. Such aging clocks generally pay little attention to the difference between females and males. However, our results advocate for a sex-specific age model. In addition, certain age-related diseases may accelerate aging process, as tumors aged 40% faster than matched normal tissues based on DNA methylome ^72^. Therefore, we speculate that constructing separate aging clocks in distinct sexes will help model the progression of age-related diseases accurately.

## Materials and Methods

### Transcriptome quantifications and data processing

We used the transcriptome data from Genotype-Tissues Expression (GTEx) project version 8.0, which contains 17,382 samples in 54 tissues (including 13 brain regions and two cell lines). We downloaded the RNA-seq bam files and the phenotype information from dbGaP (study accession: phs000424.v8.p2; table accession: pht002743.v8.p2.c1). The tissues with a small number of samples and specific to only a single sex (such as ovary and testis) were removed for a better comparison. In addition, for each donor, we combined 13 brain regions into 4 main functional regions, including hormone- or emotion-related region (amygdala, anterior cingulate cortex, and hypothalamus), movement-related region (cerebellar Hemisphere, cerebellum, and spinal cord), memory-related region (caudate, hippocampus, nucleus accumbens, putamen, and substantia nigra) and decision-related region (cortex, and frontal cortex). As a result, for the N donors that each have 13 brain regions (N×13 sample matrix), we merged the data into 4 functional regions to generate an N×4 sample matrix in the downstream analyses. In total, 16,202 samples in 35 tissues or brain regions (31 tissues and 13 brain regions) from 948 donors passed these filters for further analysis. The numbers of samples in each tissue are shown in table S1.

GE and AS quantifications were performed by Paean ^73^, a parallel computing system on the GPU-CPU platform with high computational efficiency for super-large datasets. We calculated TPM (transcripts per million) for protein-coding genes and PSI (percent of spliced-in) values for AS events, including skipped exon (SE), alternative 5′ splice site (A5SS), alternative 3′ splice site (A3SS), and retained intron (RI). Genes with low expression (average TPM < 1) were removed. We also filtered the AS events by first removing the samples with missing PSI values in more than 50% of AS events (16,182 samples left), and then filtered the AS events using the following criteria: (1) percentage of the missing PSI values in < 5% samples; (2) averaged total reads counts in spliced junctions (i.e., spliced in counts + spliced out counts) > 10; (3) non-constant PSI values across samples; (4) max(PSI) – min(PSI) > 0.05; (5) standard deviation > 0.01; (6) averaged PSI value in the range from 0.05 to 0.95; (7) TPM of the spliced genes >1. We focused on the AS events in protein-coding genes. In different tissues, the average numbers of genes and AS events for downstream analysis were 12,144 and 13, 939, respectively.

Even after removing the AS events with too many missing values, the remaining AS data were still too sparse in some samples for a reliable statistical analysis. On average there are about 0.17% AS events missing from the RNA-seq samples in different tissues. For a single RNA-seq sample in certain tissues, there were up to 15% AS events still missing (i.e., below the detecting limit). Several imputation techniques (reference-based and reference-free) have been widely used and easily implemented in multi-omics datasets ^74^, and we used the k-nearest neighbors (KNN) algorithm for missing PSI value imputation in this analysis.

### Principal component-based signal-to-variation ratio (pcSVR)

To evaluate the sex/age effect on human transcriptome, we designed a principal component-based signal-to-variation ratio (pcSVR) based on principal component analysis (PCA) and signal-to-noise ratio (SNR). PCA process could maximize the global transcriptomic variations, and SNR could calculate the distances between sex/age groups divided by the noises within each group (Equations 1-3, female *vs.* male and young *vs.* old). We selected the principal components that captured more than 80% of the global variation (other PC cutoffs capturing 50%-90% of variations were also tested as controls). The SNR of the principal component score was calculated according to the following formula:

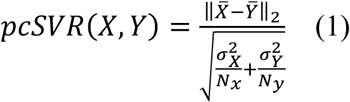

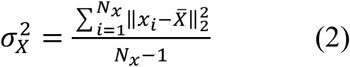

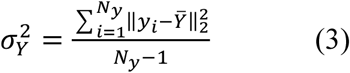

where *X* and *Y* label two groups with sample sizes *N_x_* and *N_y_* (females and males; young and old); *x_i_* and *y_x_* are the PC scores in sample *i*; and *X̅* and *Ȳ* are average PC scores across samples in each group calculated by GE or AS.

Here, to figure out whether the pcSVR is statistically significant enough, we conducted the permutation analysis and derived empirical p-values of the pcSVR. We first randomly selected subsamples (50% numbers of the minimum sample sizes of two groups) 10,000 times in each sex/age group and computed the averaged pcSVR. Then, we also randomly selected the same number of subsamples 10,000 times in each tissue regardless of the sex/age labels. The empirical p-values were calculated by the percentage of tested 10,000 pcSVR values (i.e., calculated using 50% sub-sampling irrespective of the group labels for 10,000 repeats) higher than the average pcSVR (i.e., calculated using 50% sub-sampling with the group labels for 10,000 repeats). The null hypothesis is that the two sets of pcSVR values are drawn from the same distribution. We performed all of the random sampling approaches using even probabilities in the R function ‘sample’ with the parameter prob = NULL. The tissues with more than 3 samples in each age group in females or males are included for this analysis (total 29 tissues and 4 brain regions).

To compare different approaches for empirical p-value estimation, a bootstrap with the replacement approach was also performed for null distributions. The process was repeated 10,000 times to minimize the biases of random sampling to construct a proper null distribution. Compared with the boot-strapping approach with replacement, the sub-sampling approach could also evaluate the robustness of pcSVR values to the sample size. Genes and AS events encoded by the protein-coding genes with averaged TPM > 1 were considered. To evaluate the contribution of sex chromosomes to pcSVR, we also removed the genes and AS events encoded by sex chromosomes and recalculated the pcSVR with corresponding p-values in each tissue.

### Identification of differentially expressed genes and AS events

A recent study reported that DESeq2 and edgeR have unexpectedly high false discovery rate (FDR) in analyzing large-scale datasets, as well as the inconsistency between differentially expressed genes identified by these two approaches ^43^. Linear regression models have been widely used for transcriptomic and genomic differential analysis in recent studies ^22,32,35,75^, which have the advantages of controlling all known (e.g., sex, age, race, etc.) and unknown variables (e.g., confounders and other batch effects, etc.). In each tissue, we fitted a linear regression model including age, sex, and age-by-sex interactions as covariates, as well as other confounding factors. The following linear regression was used:

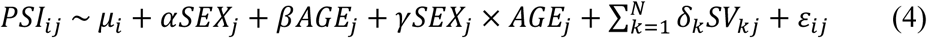

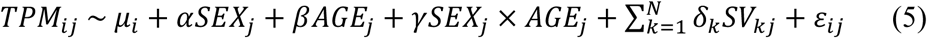

where *PSI_ij_* is the normalized PSI value for AS event *i* in sample *j*; *⍺*, *β*, *γ* are the coefficients of sex, age, and age-by-sex interactions, respectively; *μ_i_* is the regression intercept; *ɛ_ij_* is the error term; and *k* is the number of surrogate variables (SVs) in *N* total SVs for sample *j* with the coefficients *δ_k_*. SVs are estimated using SVA (Surrogate Variable Analysis)^76^. For differential GE analysis, we used the same linear regression model for analyzing differential GE by replacing PSI with TPM in the formula (Equation 5). To deal with the different orders of magnitude between PSI and TPM values, we normalized the PSI/TPM of each gene/AS event across samples using a scaling method to normalize the distance from the mean PSI/TPM across all samples.

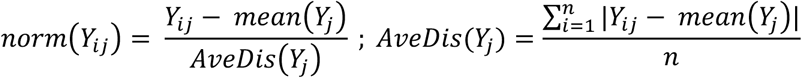

Here, *Y_ij_* indicates the PSI or TPM of the gene/AS event *i* in sample *j*, and the *AveDis*(*Y_j_*) indicates the average distance from the mean PSI/TPM across all samples.

We defined sex-differential or age-differential genes with the cutoffs of p-value < 0.05 for the α or β coefficients, and foldchange >1.5. Sex-differential or age-differential AS events were defined with the cutoffs of p-value < 0.05 for the α or β coefficients, and |ΔPSI|>0.05. We performed the differential analysis in each gene or AS event independently. The *γ* significantly larger or smaller than 0 indicates significant age-by-sex interaction, which suggests that the values of TPM and PSI are not only determined by sex or age independently but also affected by the age at a specific sex or sex at a specific age.

To evaluate whether the genetic effects could be controlled by SVs, we calculated the correlation between SVs and donors’ ethnicity using Point Biserial Correlation which measures the association between continuous variables and dichotomous variables. Also, the Spearman’s correlation was used to calculate the correlation between SVs and the top5 principal components as judged by whole genome sequencing data (WGS). The ethnicity information was extracted from phenotype data. The results of principal components calculated by the WGS dataset were downloaded from GTEx portal.

Permutation analysis was used to address the false discovery issue of this linear regression model in the differential analysis. We first generated the permuted data by randomly shuffling the group labels (sex or age) with the same sample sizes as the original group labels. The differential genes/AS events (DEGs/DASs) were identified based on each generated permutated phenotype data. This process was performed 1,000 times and we used two strategies to examine the robustness of our cutoffs. We first defined the fractions of original DEGs/DASs which are also identified in each permutation iteration. Meanwhile, the FDR is defined as how many permutations incorrectly identified the specific DEG/DAS across 1,000 permutations.

### Evaluation of the effect of the sample sizes in differential GE and AS analysis

The effects of GTEx sample sizes on the numbers of differential genes/AS events were evaluated by a linear regression model by the following formula:

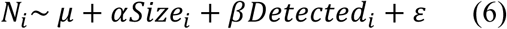

where *N* is the number of sex-differential or age-differential genes/AS events in tissue *i*; *Size* is the number of selected samples; *Detected* is the detected numbers of genes/AS events after filtering in preprocessing procedures; *⍺*, *β* are the coefficients of the sample sizes and numbers of detected genes/AS events, respectively; *μ* is the regression intercept; and *ɛ* is the error term.

### Identification of sex-stratified age-associated genes/AS events

To identify the genes/AS events affected by age in separate sexes, we also conducted differential analysis by fitting the following linear regression model in females and males, respectively:

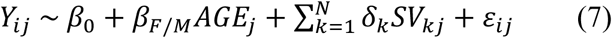

where *Y_ij_* is the normalized TPM or PSI values of gene/AS event *i* in sample *j*; *β_F/M_* is the coefficient of age in females or males; *β*_0_ is the regression intercept; *ɛ_ij_* is the error term; and *k* is the number of SVs in *N* total SVs for sample *j* with the coefficients *δ_k_* evaluated in each sex. Age-associated genes or AS events were defined according to a significant age effect in different sexes (βM, βF, with cutoffs of p<0.05; GE fold change>1.5 or AS |ΔPSI|>0.05). The biases in different sexes also reflected the functional interactions between sex and age. This approach not only could indicate whether there is a sex-by-age interaction, but also define the direction of sex-by-age interaction, that is, how females/males affect GE/AS during aging differently. In addition, ΔPSI and Foldchange have also been used for filtering, which are the important thresholds to evaluate the degree of changes. Thus, we further defined the female- or male-specific age-associated genes/AS events using this sex-stratified analysis, and classified these age-associated genes/AS events into sex-biased age-associated genes/AS events (i.e., age-associated genes or AS events specific to one sex) or sex-ubiquitous age-associated genes/AS events (i.e., age-associated genes or AS events in both sexes).

### Functional enrichment and Disease Ontology analysis

Potential functions of sBASEs were expounded based on C5 collections (ontology gene sets) in Molecular Signatures Database (MSigDB) and Disease Ontology. We carried out these processes using Bioconductor R packages *msigdbr* ^77^, *clusterProfiler* ^78^, and *DOSE* ^79^, with the significance threshold p-value <0.05. The GO terms with the significances specific in only one sex per tissue were selected for downstream analysis. In the enrichment heatmap of brain regions (Fig. 4A), we plotted the GO terms with significance in at least 3 brain regions. Because the enrichment heatmap across all tissues are unrealistically large, we plotted the terms within at least 15 tissues and the tissues/sex with significance in at least 9 significant GO terms (fig. S8). The entire results of the enrichment analysis were shown in table S4.

### Processing of Alzheimer’s disease datasets and construction of AD prediction model

The AD samples of human prefrontal cortex were selected from GEO datasets (ID: GSE174367) containing 375 samples of single-nucleus RNA-seq, single-nucleus ATAC-seq, and bulk RNA-seq. We filtered the datasets by selecting the RNA-seq samples from the Frontal Cortex brain region (Library Source: ‘TRANSCRIPTOMIC’; Brain region: ‘FC’). Next, we selected the samples with the isolation approach ‘total RNA isolation’. Due to the sequencing depth on the splicing junction, we filtered out the samples lower than 1Gb Bytes (SRR14514282, SRR14514349, SRR14514308, and SRR14514275). The remaining 91 samples were selected for GE and AS quantification, including 45 AD samples and 46 control samples from 47 male and 44 female donors (Female:Male≈1:1; AD:Control≈1:1) with similar age distribution between AD and control group. Quality control and adaptor trimming were performed using FastQC ^80^ (version 0.11.9) and Trim Galore (version 0.6.6). Bowtie2 ^81^ (version 2.4.2) was used to remove reads mapping to ribosomal RNA. Then, clean reads were mapped to human reference (GRCh38) using STAR ^82^ (version 2.7.6). We also used pre-mapped RNA-seq data from ROSMAP containing the RNA-seq samples from the bulk sequencing of DLPFC. We further filtered the data to match the number of both sexes and the age distribution between AD and control groups, resulting the 215 AD and 56 control samples. The quantifications of GE and AS were performed by Paean ^73^.

To identify AD-related genes/AS events, we fitted a linear regression model with AD diagnosis (categorical outcomes) and surrogate variables calculated by SVA ^76^. AD-related genes/AS events were defined whose coefficients of the ‘diagnosis’ term significantly deviated to zero. To compare the correlation coefficients that evaluated the correlation between AD-associated and age-associated AS changes in males and females independently, we performed Fisher’s z-transformation on Pearson correlation coefficients using R packages *cocor* ^83^.

To capture the key age-associated splicing changes in males and females that particularly involved in AD pathogenesis, we focused on in the frontal cortex that relevant to AD and further use the sBASEs for AD prediction. We specifically used sBASEs that exhibited specific sex-biased changes in splicing associated with aging (315 female-biased sBASEs and 208 female-biased sBASEs). This subset of sBASEs was chosen in terms of those that could also be detected in the ROSMAP AD dataset due to different sequencing depths or technical biases across datasets. These sBASEs were further input to a prediction model with the feature selection algorithm and then evaluated their contributions. We randomly divided 90% samples for training sets and 10% for test sets and repeated for 100 iterations. In each iteration, we performed a 5-fold cross-validation with the feature selection approach called Recursive Feature Elimination (RFE) via R packages *caret* ^84^. Model performance was evaluated by the area under the curve (AUC). Random forest (RF) and support vector machine (SVM) with a linear kernel were used for classification (Fig. 4D and fig. S9D). To evaluate the final selected features that crucially contributed to AD prediction, we calculated the importance of each feature in each iteration (i.e., mean decrease accuracy, MDA) and then ranked the features by the averaged MDAs across 100 iterations. As negative controls, the sex-stratified model predicted AD patients in females and males separately using randomly selected AS events, and the merge-sexes model predicted AD patients in all samples regardless of sex labels using sBASEs.

### Transcriptome-wide association analysis with AD pathology

We used a linear regression model to identify the associations between AS events and AD pathology. The pathology includes tangles and plaque stages with continuous outcomes. Age was included as one of the covariates. This association analysis was performed for each sex.

### Construction of AS regulatory network

Splicing-related genes were extracted from the GO_RNA_SPLICING term based on MsigDB (c5.all.v7.4.symbols.gmt). Networks between age-associated splicing factors (SFs) and sBASEs in each sex were constructed with the following three steps. Firstly, we examined the significant correlations between the expression of SFs and splicing of AS events during aging. We conducted spearman’s correlation test between TPMs of SFs and PSIs of AS events during aging using the threshold p-value < 0.05.

Secondly, we chose the AS events significantly perturbed by RBP knockdown for downstream analysis. RBP shRNA RNA-seq bam files of 200 RBPs in HepG2 and K562 were downloaded from ENCODE Consortium. The TPM and PSI values were also quantified by Paean ^73^. Differential analysis between control and shRNA groups was performed by a linear regression model:

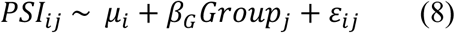

where *Group_j_* represents a categorical variable labeled by “Control” or “shRNA”. AS events significantly perturbed by RBP knockdown were defined by the estimated coefficient *β_G_* significantly deviated from zero.

Thirdly, we identified the AS events with SF binding motifs around the alternative splice sites (SS). We used DeepBind ^85^ to predict the specific binding scores of RBPs within the upstream to downstream region adjacent to the alternative SS. BEDTools getfasta ^86^ extracted sequences from specific genomic regions in a .bed6 file. We first subdivided the sequences into multiple subsequences (window = 40nt; step =1nt) and calculated the binding scores with the parameter ‘average’ of each subsequence. To identify whether there were binding signals in the upstream to downstream 300nt region adjacent to the alternative splice sites, we rearranged the binding scores in each region into small bins (width = 20), and tested the median differences between the bins with maximum and minimum averaged binding scores by Wilcoxon rank-sum test. The AS events whose maximum binding scores > 1 and p-value < 0.05 were reserved. Taken together, we required the regulatory connections between the RBPs and AS events that pass all the individual criteria as judged by all three types of experiments with different techniques. The networks between age-associated SFs and sBASEs were constructed and figured by Cytoscape ^87^.

### Regulation of hormone receptors to the target genes

To examine the estrogen regulation using the RNA-seq dataset, we downloaded the RNA-seq datasets from MCF-7 cell line stimulated by estrogen receptor agonist (propyl pyrazole triol, PPT) in three biological replicates (GEO ID: GSE203058) ^48^. Additional datasets (ID: GSE89888) are downloaded to confirm our findings in engineered MCF-7 cell lines ^49^ with wildtype and two constitutively active mutants (Y537S, and D538G) of ESR1. These cells were treated with 1nM estradiol (E2) or vehicle control (veh) for 24 hours before collecting the total RNAs and subjected to transcriptome profiling by RNA-seq. We analyzed these data using the same pipelines in earlier Alzheimer’s disease datasets (Fig. 4). The GE changes under estrogen treatments were evaluated in a linear regression model using the p-values of the coefficients deviated from zero. To confirm the expression changes of splicing factors in other cells or tissues, we downloaded the FPKM table from GSE86609 that analyzed the ER knockout mice with estrogen treatments in mouse brain arcuate nucleus (ARCs) ^50^.

To further identify the regulation of the estrogen and androgen using ChIP-seq dataset (Fig. 5E and Fig. 6H), we evaluated the binding scores of ESR1 and AR to the promoter regions of the target genes (i.e., +/− 1kb of the transcription start sites, TSS). The binding score of each target gene (i.e., −10*log10[MACS Q-value] from ChIP-seq analysis) was calculated by the ChIP-Atlas ^88^, an integrative database of public ChIP-seq data. The ratios in Fig. 5E were calculated by the binding scores of each SFs divided by the median binding score of age-associated SFs common in both sexes. Additionally, the ESR1 ChIP-seq peak files in WT mice treated by estrogen were also downloaded from the Gene Expression Omnibus (ID: GSE36455) ^51^. The peaks were annotated using ChIPSeeker ^89^.

### Time series and breakpoint analysis

Autoregressive Integrated Moving Average (ARIMA) model is one of the linear models for forecasting chronological trends. For non-stationary data, *auto.arima*() function from R package *forecast* conducts a stepwise procedure to search for the best model using the smallest Akaike Information Criterion (AIC). This procedure is a variation of the Hyndman-Khandakar algorithm ^52^.

We first corrected the TPM/PSI values for each gene/AS event by removing the unknown confounding factors except for the sex and age effect *via* R package *limma*, and averaged the values of the samples with the same sex and age. Then, we applied ARIMA models to figure out genes with potential chronological trends during aging in each sex. In brief, the input data was first transformed into z-ordered objects (R package *zoo*) and the best model was stepwise optimized by AIC (*auto.arima* function from R package *Forecast*). We filtered out the genes/AS events with the significant trend for further analysis by non-seasonal difference order higher than zero (D>0) and the sum of AR or MA coefficient higher than zero (AR+MA>0).

Furthermore, we calculated the rate of change at each age point and estimated the breakpoints during aging. The rate of change at each age *i* was evaluated by the differences between bins in (*i* − *w*) ∼ *i* and *i* ∼ (*i* + *w*) in the following steps. *w* indicated a series of window spans from 5 to 15 years old. The p-values were first tested by MANOVA of three PCs after reducing dimensionality in the PCA process and then smoothed by LOESS regression with the smoothing bandwidth in the range from 0.25 to 0.75 (with interval 0.05). The p-values of each window span were integrated by average -log10 transformed p-values and smoothed by LOESS regressions with bandwidth = 0.5. The final rate of changes at each age point was defined by the averaged values through all window spans calculated in the previous step. Moreover, we determined the breakpoints as the age points at the global maximums, as well as the local maximums whose distance to the nearest minimum relative to the global maximum was more than 10%. This model was conducted based on the approach from a previously published study ^16^ (URL: https://github.com/UcarLab/SexDimorphismNatureCommunications).

### Identification of aging-modulated genes (AMGs)

We performed the following steps to identify the AMGs that serve as the key contributors to molecular aging (Fig. 6D). We randomly trimmed 20% of chronological genes 200 times and repeated breakpoint analysis in each iteration. Wilcoxon signed-rank test was used to compare the changing rate after removal with that calculated before removal. Using a p-value < 0.05, we obtained the set of genes that could significantly alter the aging rate. The AMGs were defined as the set of genes that were significantly enriched from the 200 random samplings (as judged by the Fisher exact test using a p-value < 0.05). The tissues with more than 10 chronological genes were selected for AMGs identification process. Functional enrichment of sex-biased AMGs was based on MSigDB, and the network between GO terms and genes was constructed by Cytoscape ^87^. GE patterns during aging were shown by the R package *pheatmap*.

Our results observed clusters of AMGs common in both sexes in whole blood tissue with sex-specific changing patterns (i.e., some genes increased first at young ages and then decreased during male aging, whereas these genes monotonically decreased during female aging). To identify AMGs with this expression pattern, we divided the age points into 9 age windows (e.g., 20-25, 26-30, 31-35, etc.) and averaged the ARIMA-fitted expression levels of the samples within each age window. The AMGs increased at young ages were defined by the mean of fitted expression levels at 31-40 larger than at 20-30.

## Acknowledgments

The authors want to thank Dr. Yue Hu and Jiefu Li in Wang Lab for their discussions and comments. We also thank Prof. Guoqing Zhang in SINH for his assistance with public data access and Ruijie Yao for helping download RBP shRNA-seq bam files from ENCODE Consortium and providing Paean scripts for multi-processing on the CPU-GPU platform.

## Author contributions

S.W. and Z.W. conceived and designed the study. S.W. analyzed the data and conducted the statistical analysis. S.W. and Z.W. wrote the manuscript. D.D. X.L. provided key suggestions for study design and statistical analysis. D.D. and X.L. downloaded the GTEx RNA-seq and phenotype data.

Conceptualization: SW, ZW

Methodology: SW, DD

Investigation: SW

Visualization: SW

Supervision: XL, ZW

Writing—original draft: SW, ZW

Writing—review & editing: SW, DD, XL, ZW

## Data and materials availability

GTEx data were obtained through dbGaP (accession number phs000424.v8.p2). The source RNA-seq data (*.fastq) of AD and control samples were obtained from Gene Expression Omnibus (accession number GSE174367). The RNA-seq and ChIP-seq data with estrogen treatment in ER mutant and wild type cells were downloaded from Gene Expression Omnibus (accession number GSE203058, GSE89888, GSE86609, and GSE36455). Pre-mapped RNA-seq data (*.bam) of AD and control samples were obtained from ROSMAP (accession number syn8540863). shRNA-seq data of 200 RBPs in HepG2 and K562 cell lines were downloaded from ENCODE consortium (https://www.encodeproject.org). Molecular Signatures Database, (http://software.broadinstitute.org/gsea/msigdb/collections.jsp). The source code is available at https://github.com/rnasys/Sex-dimorphic-aging and from Zenodo https://doi.org/10.5281/zenodo.10970995. All data are available in the main text or the supplementary materials.

---

**fig. S1.**
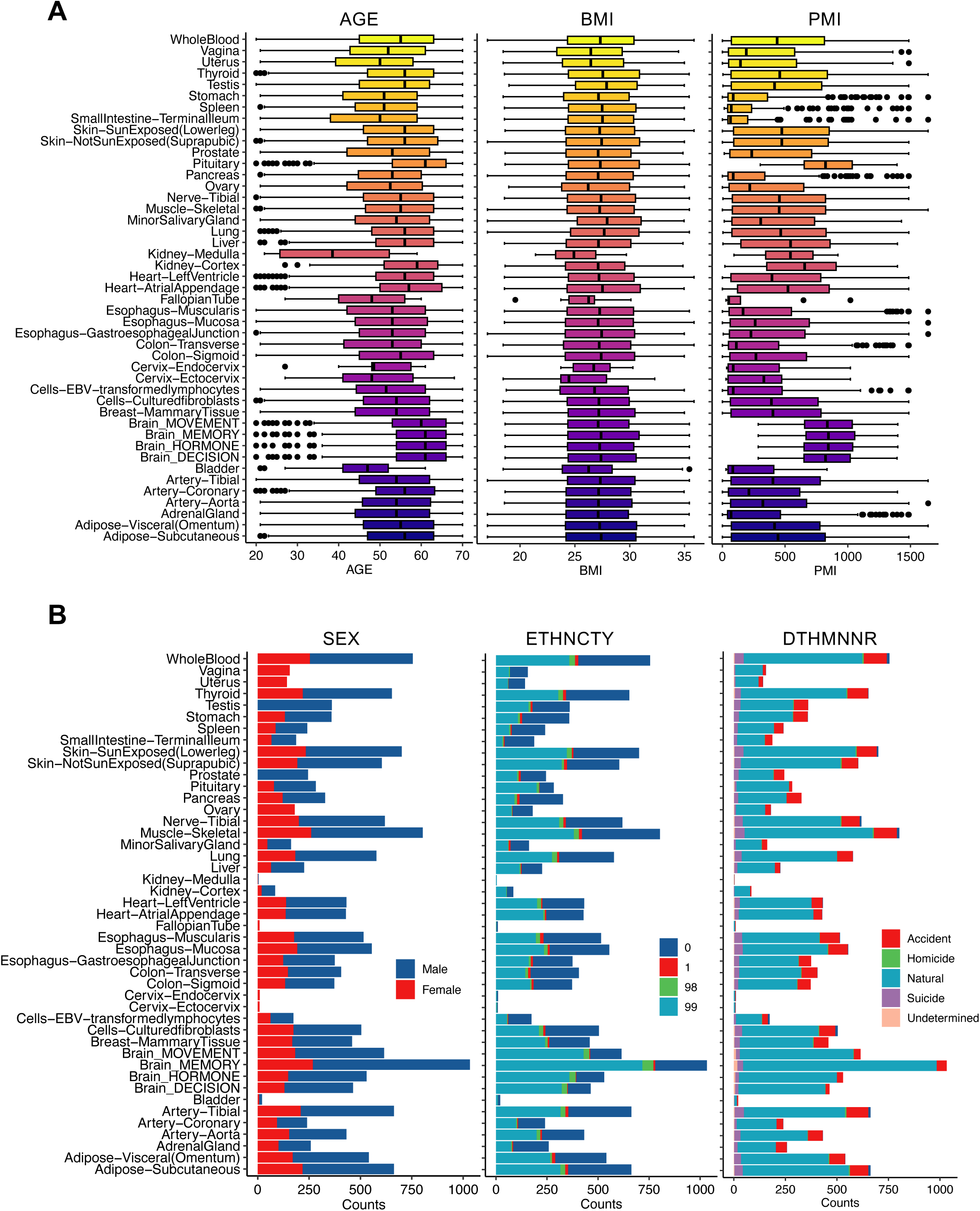
Sample phenotype in GTEx v8 cohort.

**fig. S2.**
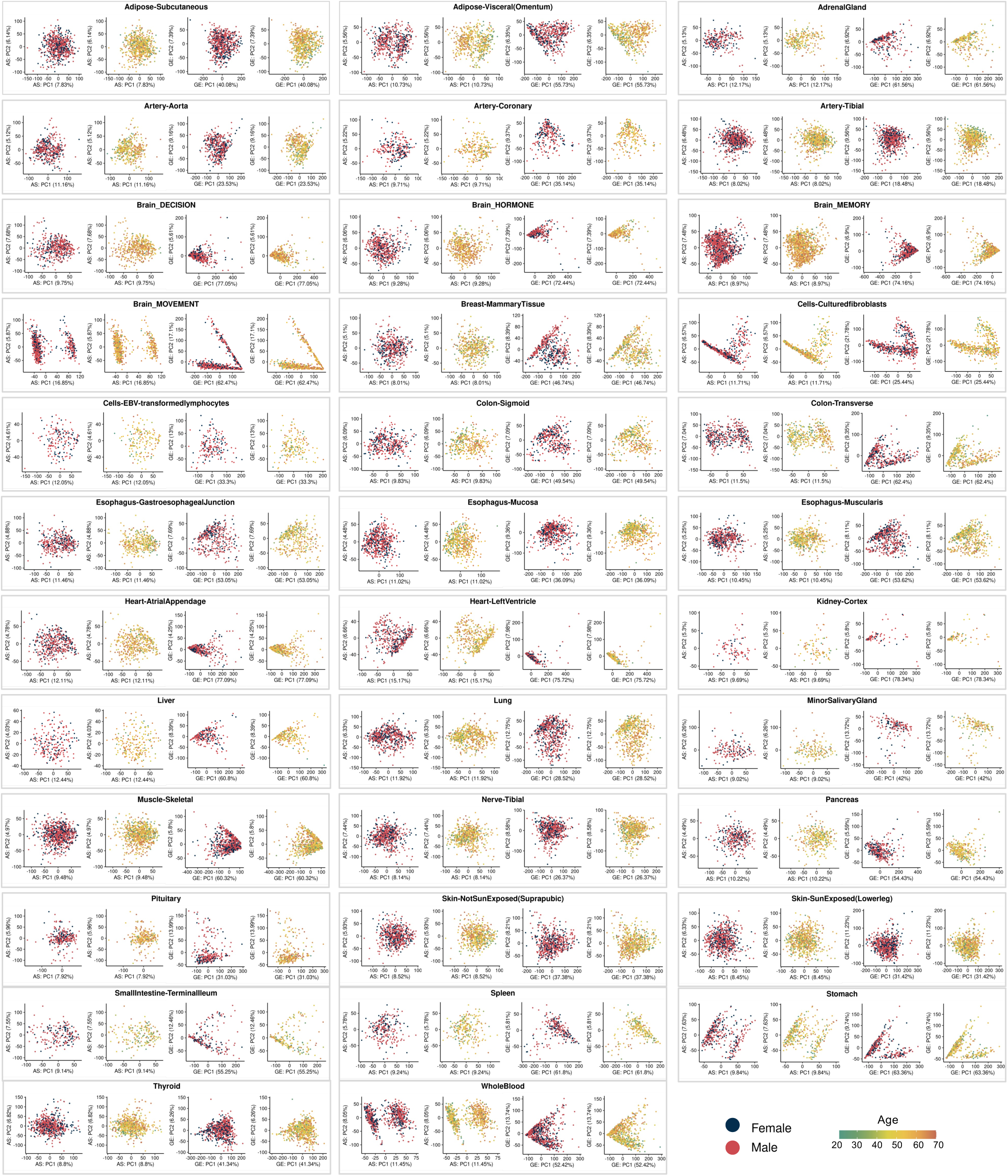
PCA on GE and AS profile.

**fig. S3.**
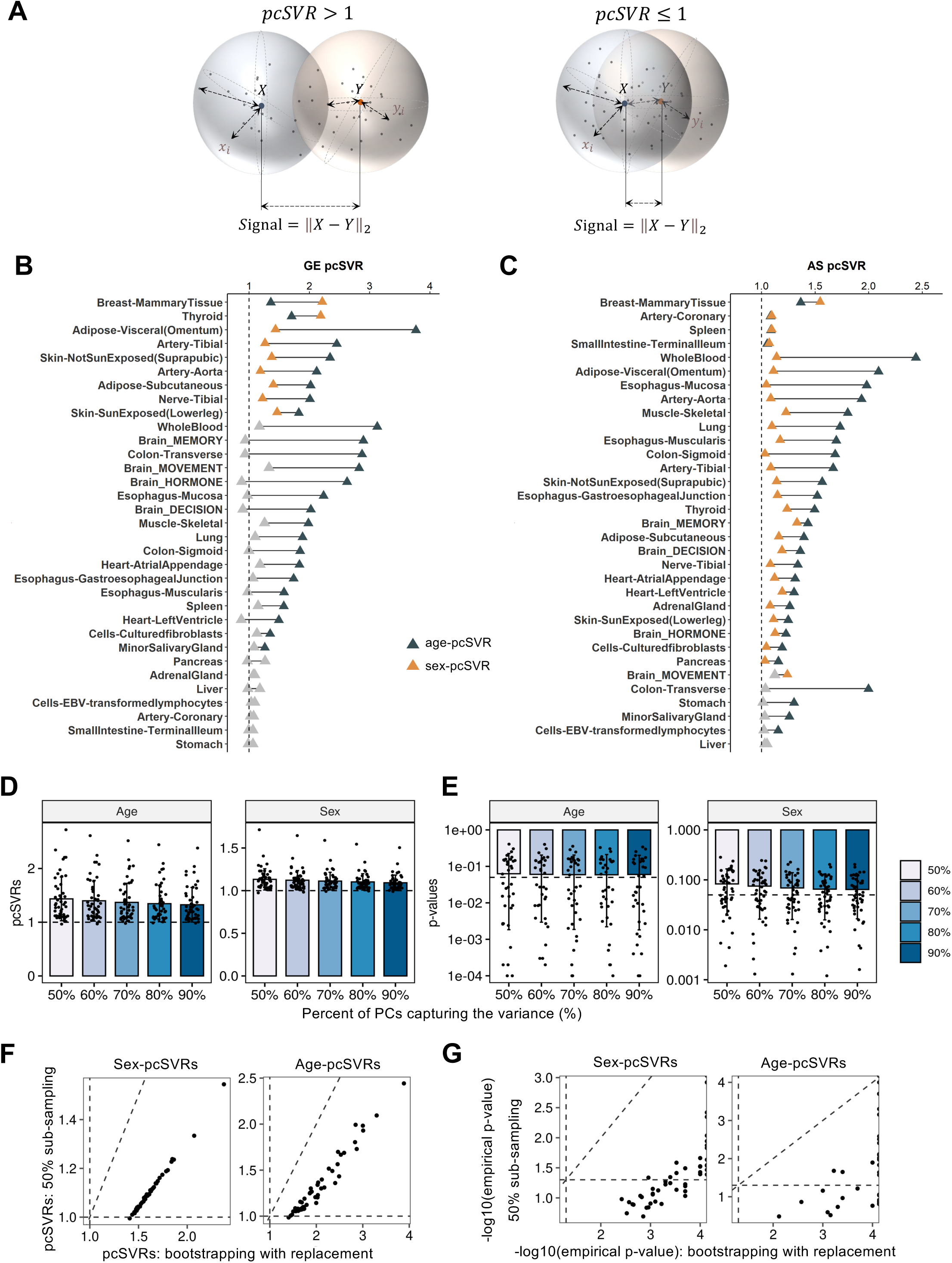
pcSVR calculation and robustness.

**fig. S4.**
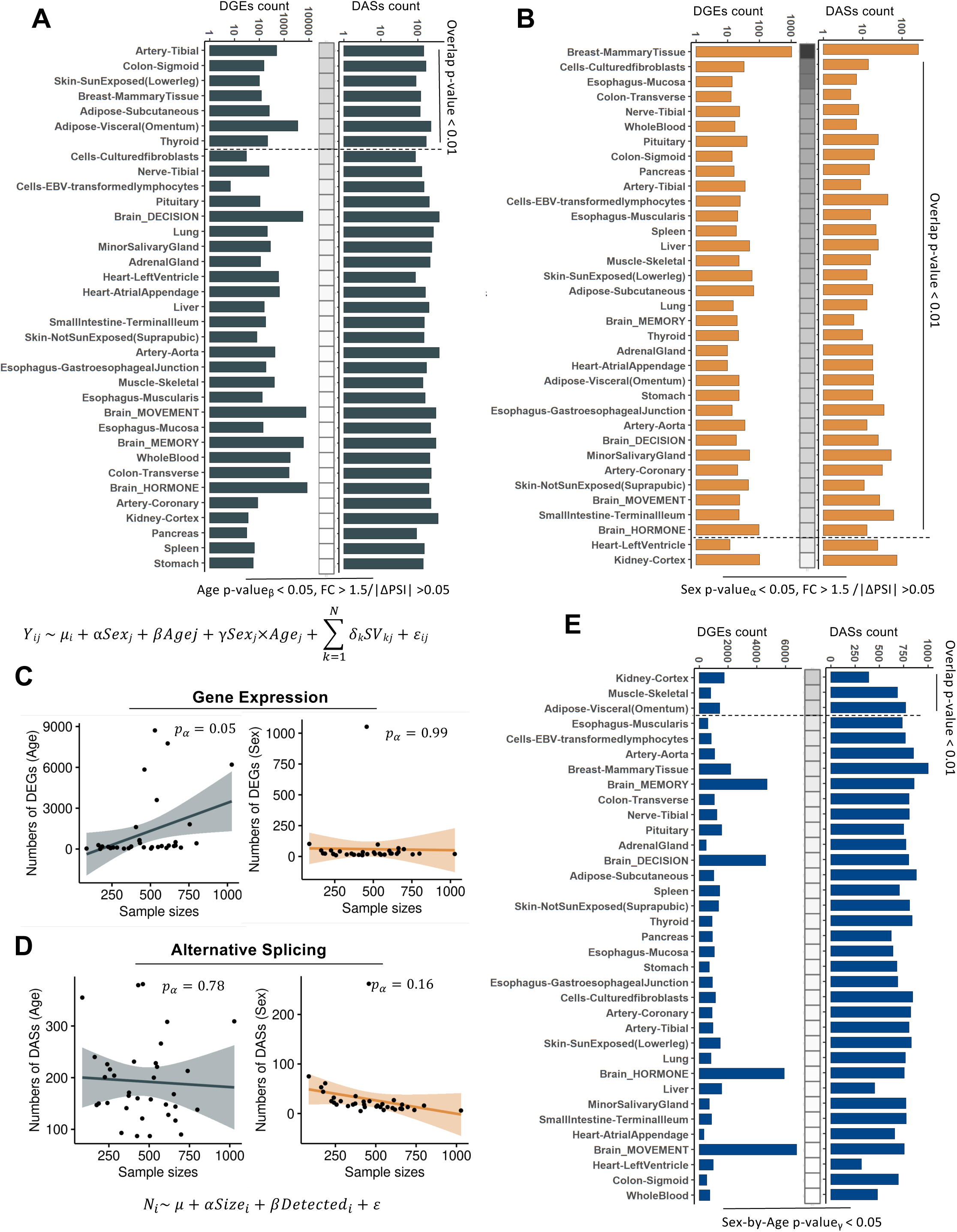
Differential GE and AS analysis to identify sex/age-differential genes and AS events.

**fig. S5.**
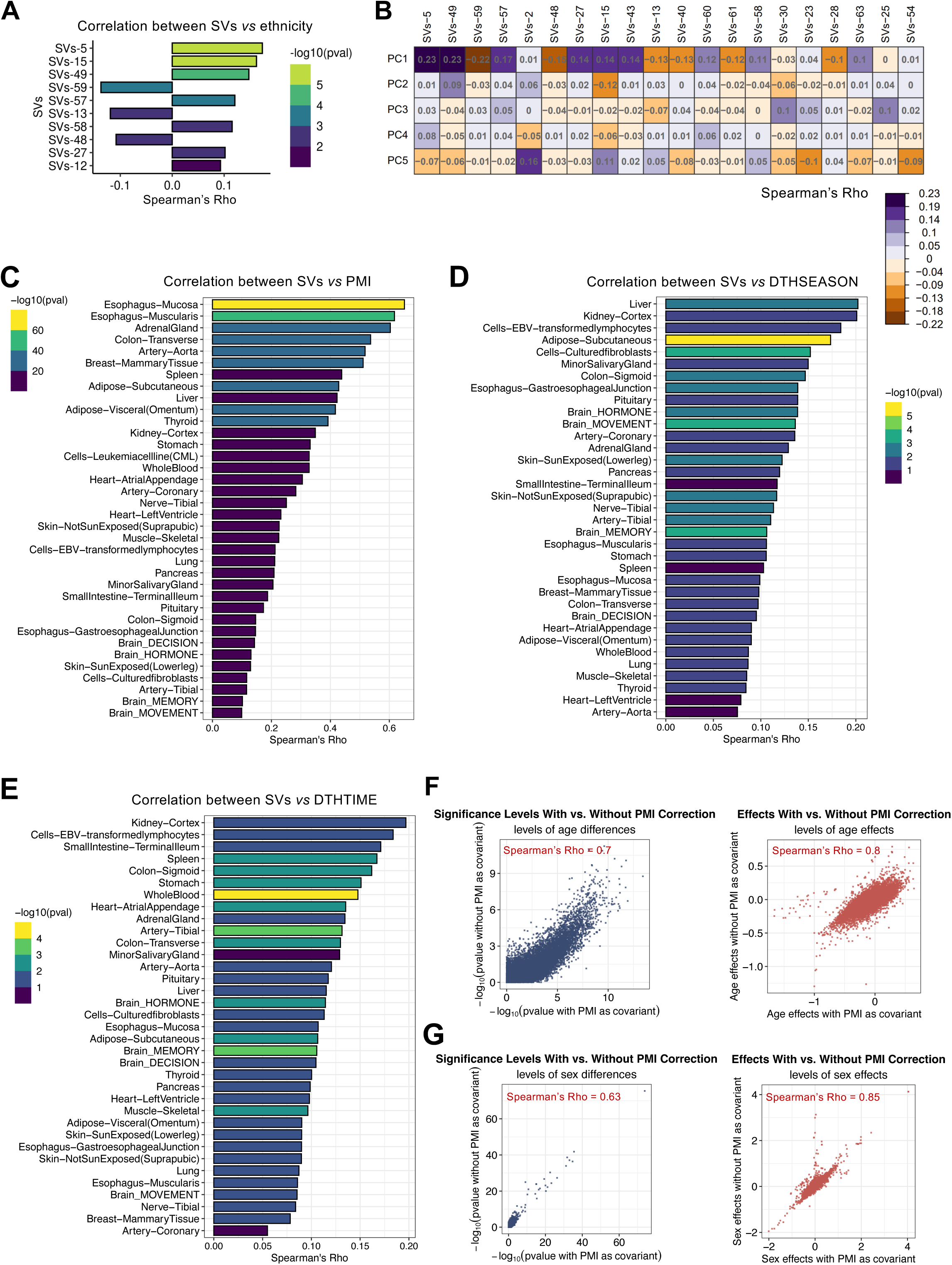
The evaluation of the confounding factors in the linear regression model.

**fig. S6.**
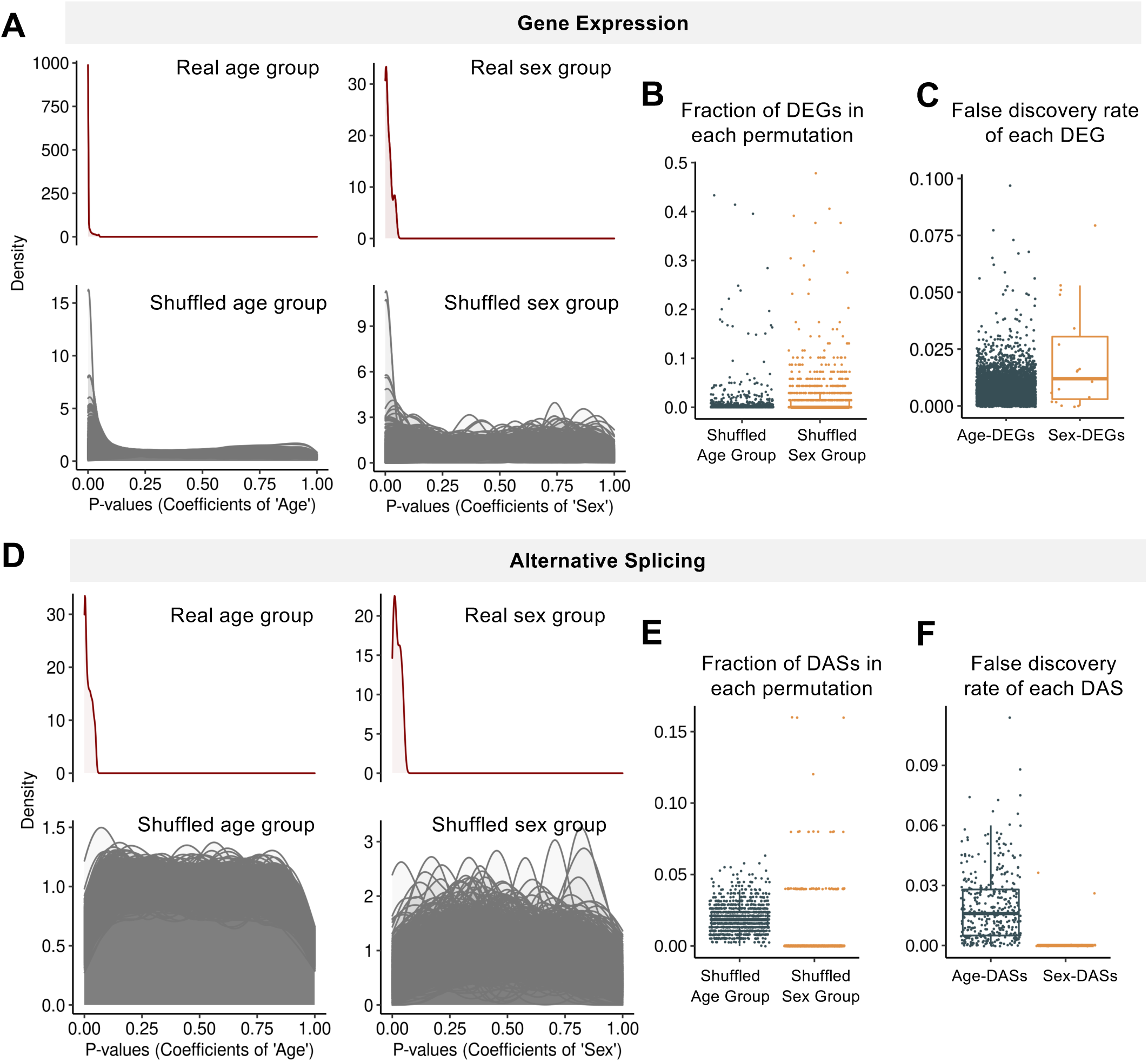
Permutation analysis of the linear regression model in differential analysis.

**fig. S7.**
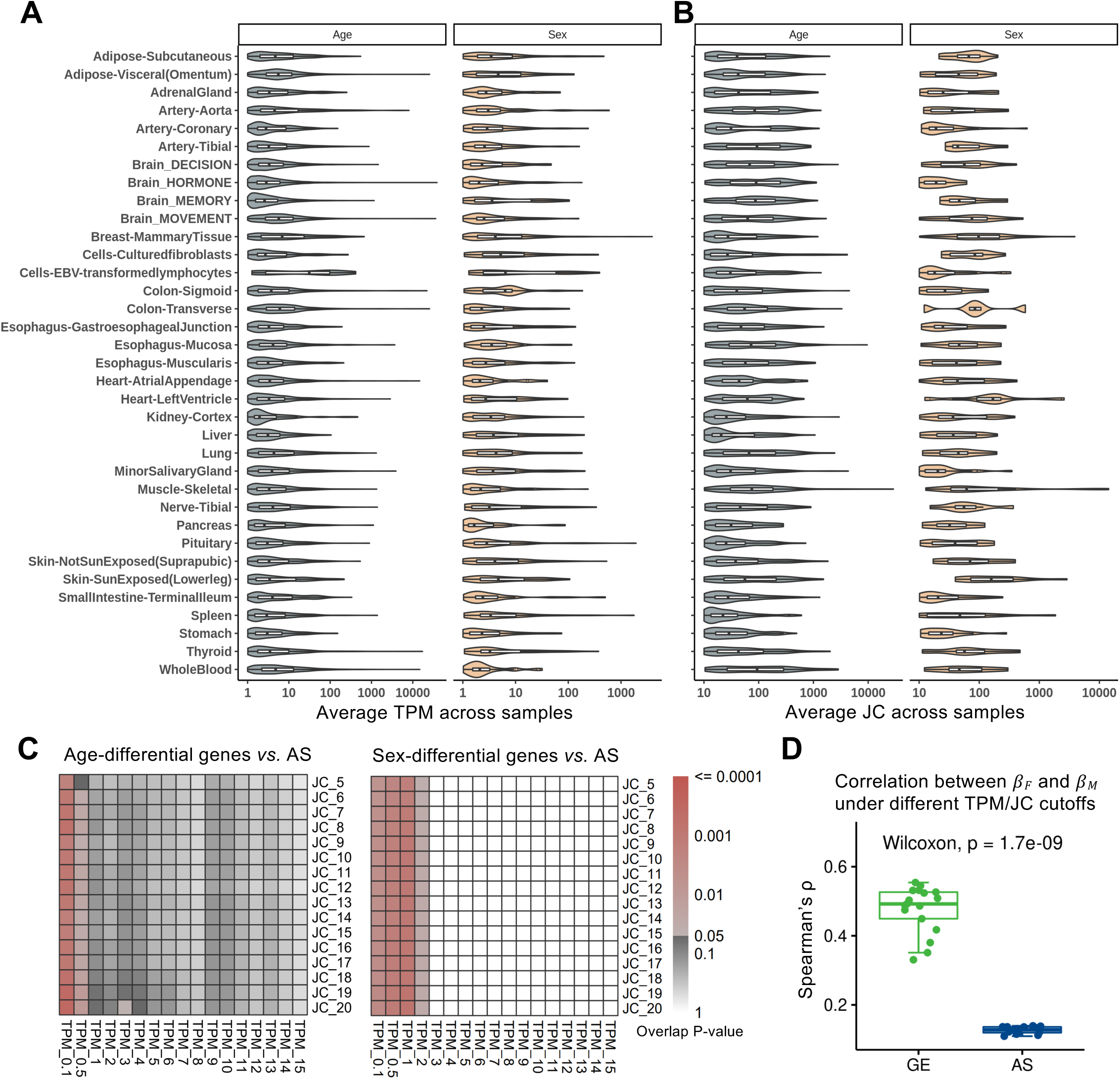
Robustness of GE and AS cutoffs.

**fig. S8.**
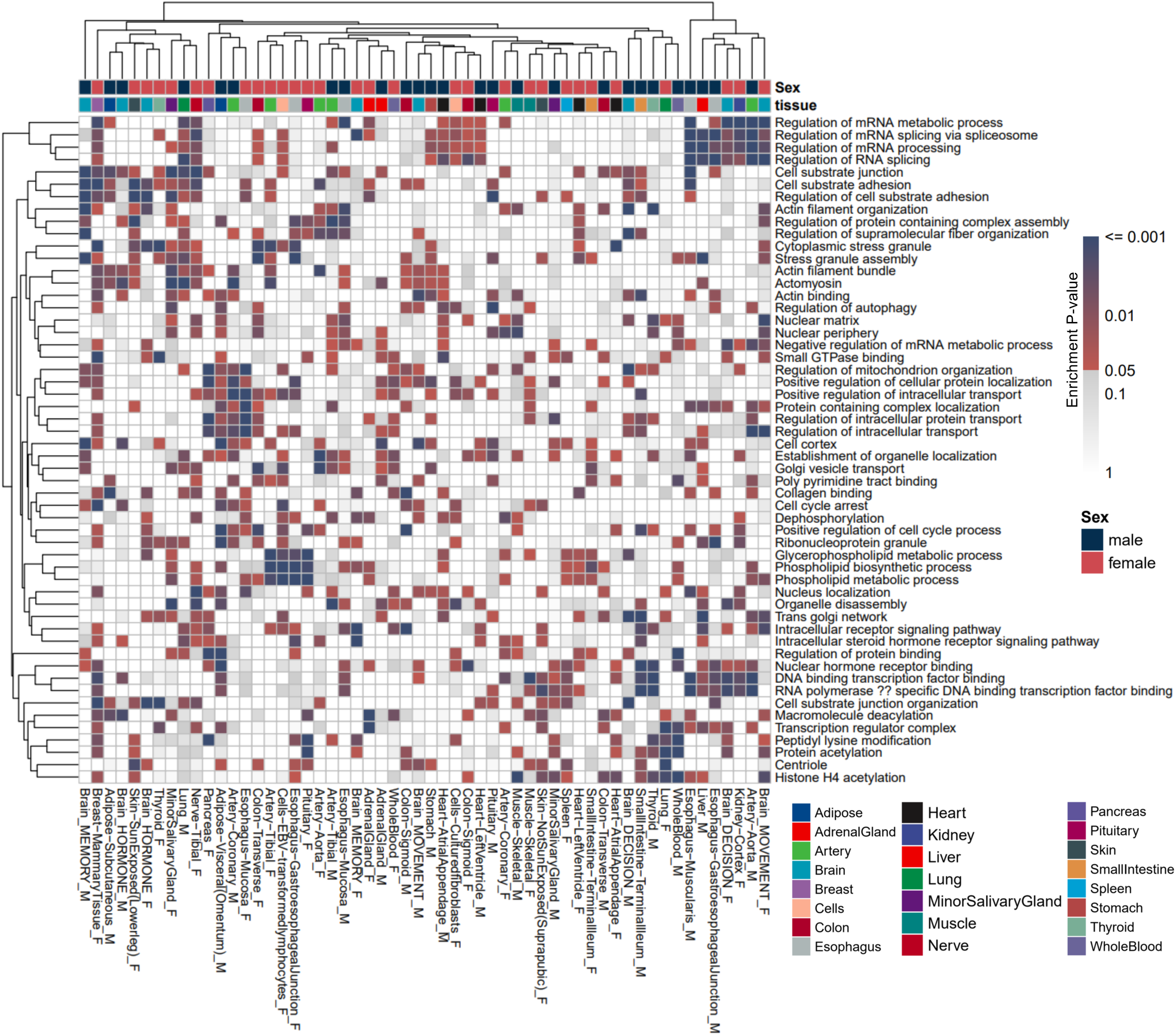
Functional enrichment of sBASEs across tissues.

**fig. S9.**
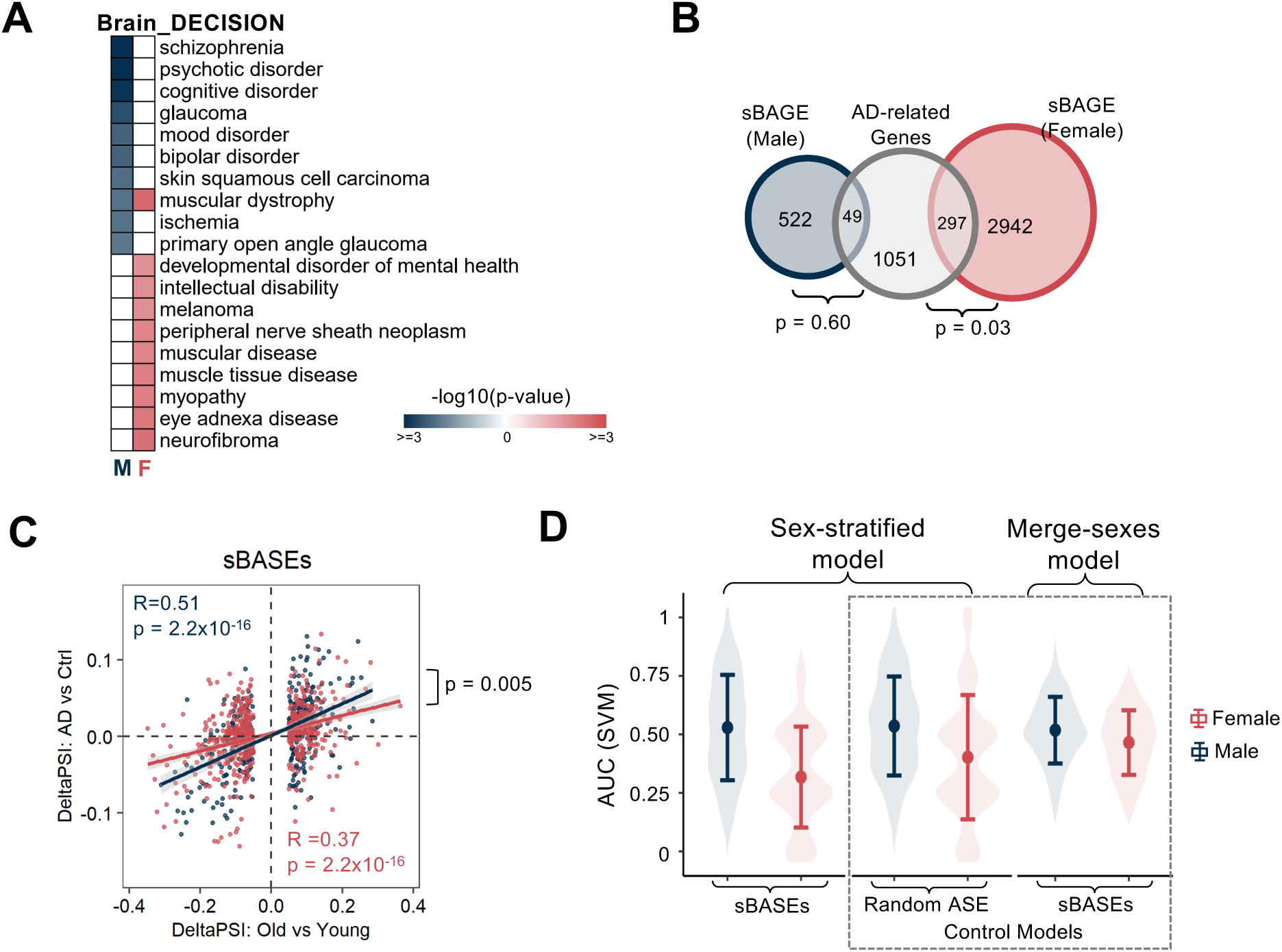
Sex-biased associations between sBASEs and diseases in decisionrelation brain region.

**fig. S10.**
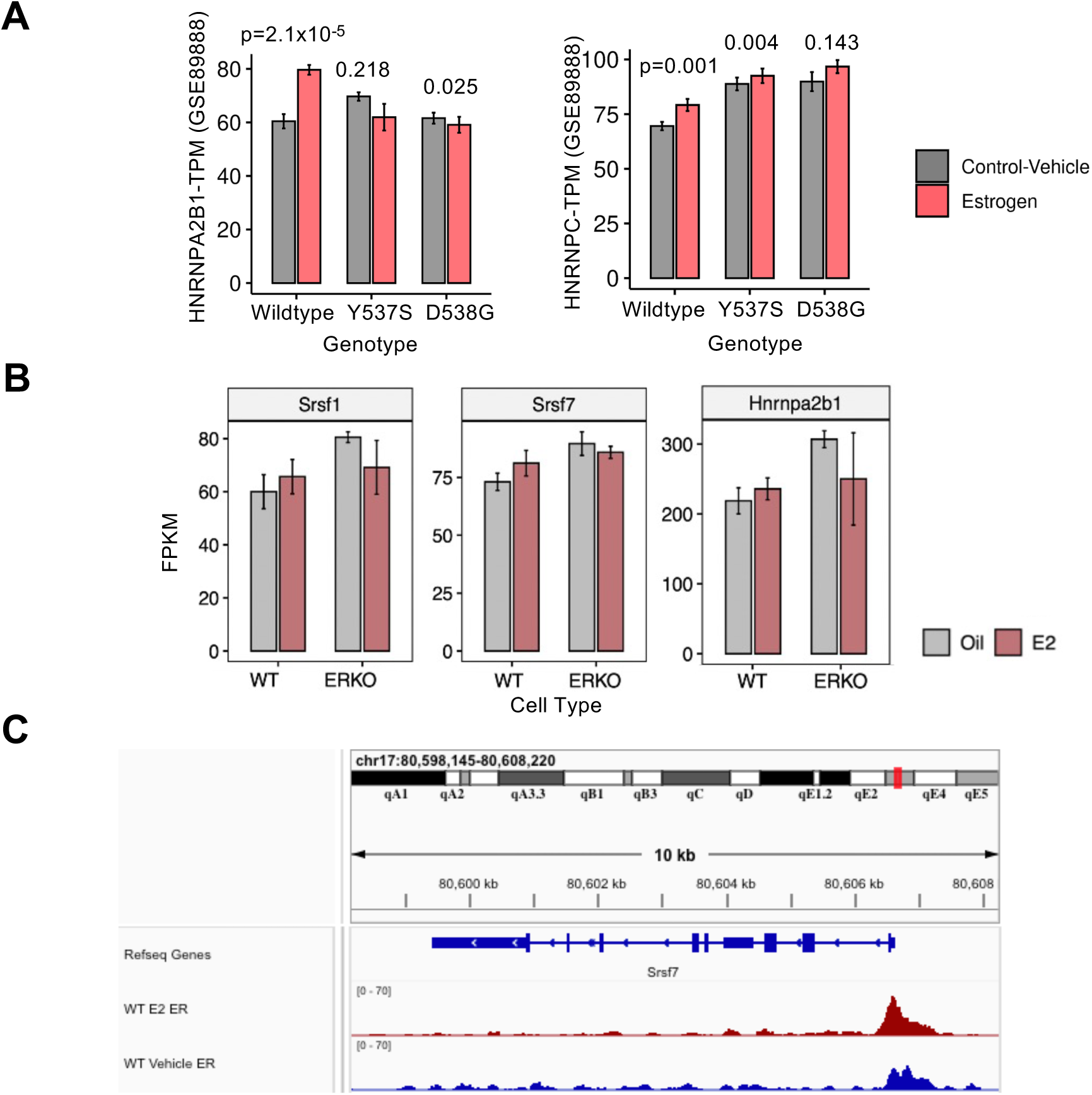
Gene expression regulation of sex-biased age-associated splicing factors by estrogen via ESR1 in multiple datasets.

**fig. S11.**
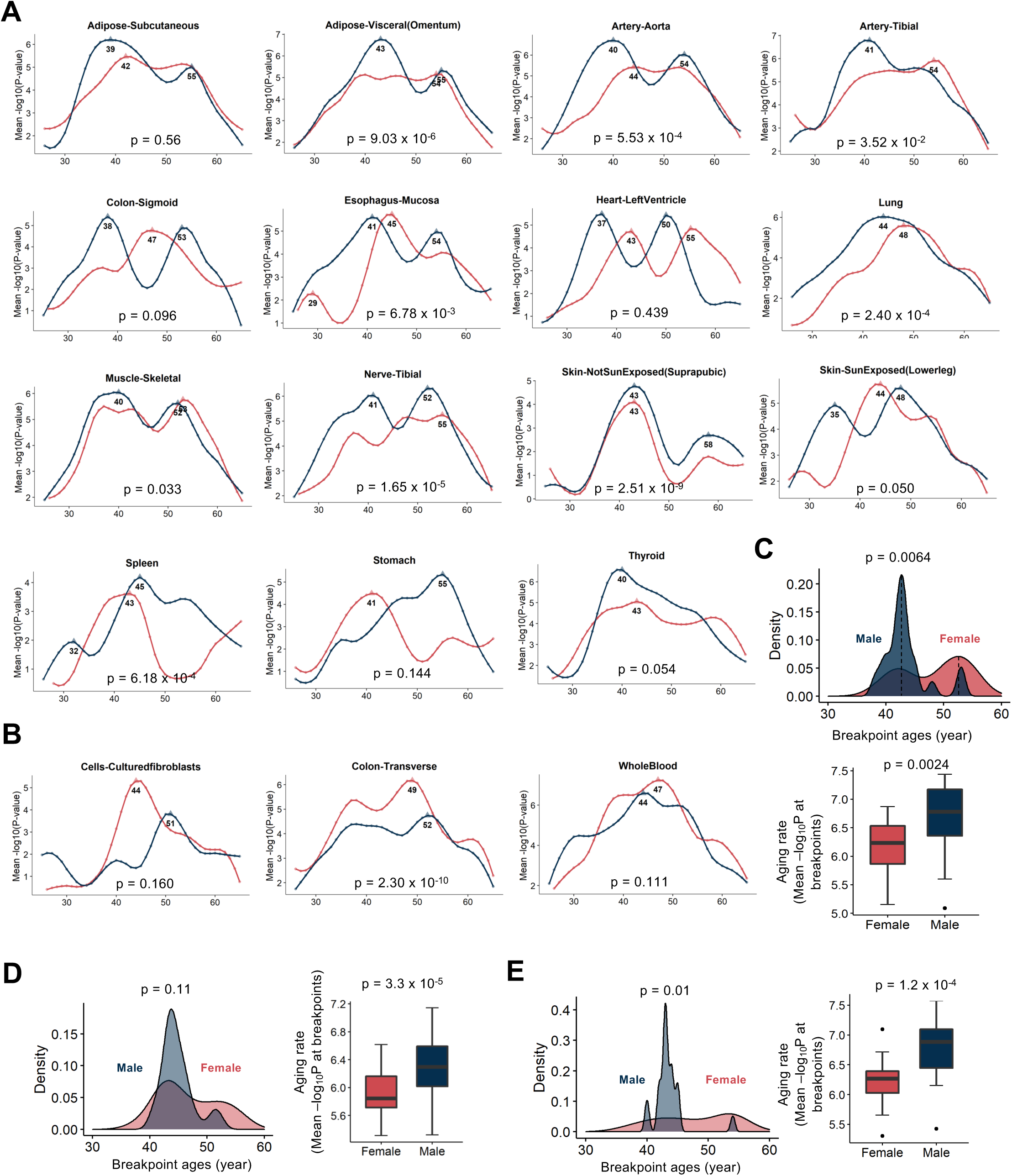
Breakpoint analysis across multiple tissues.

**fig. S12.**
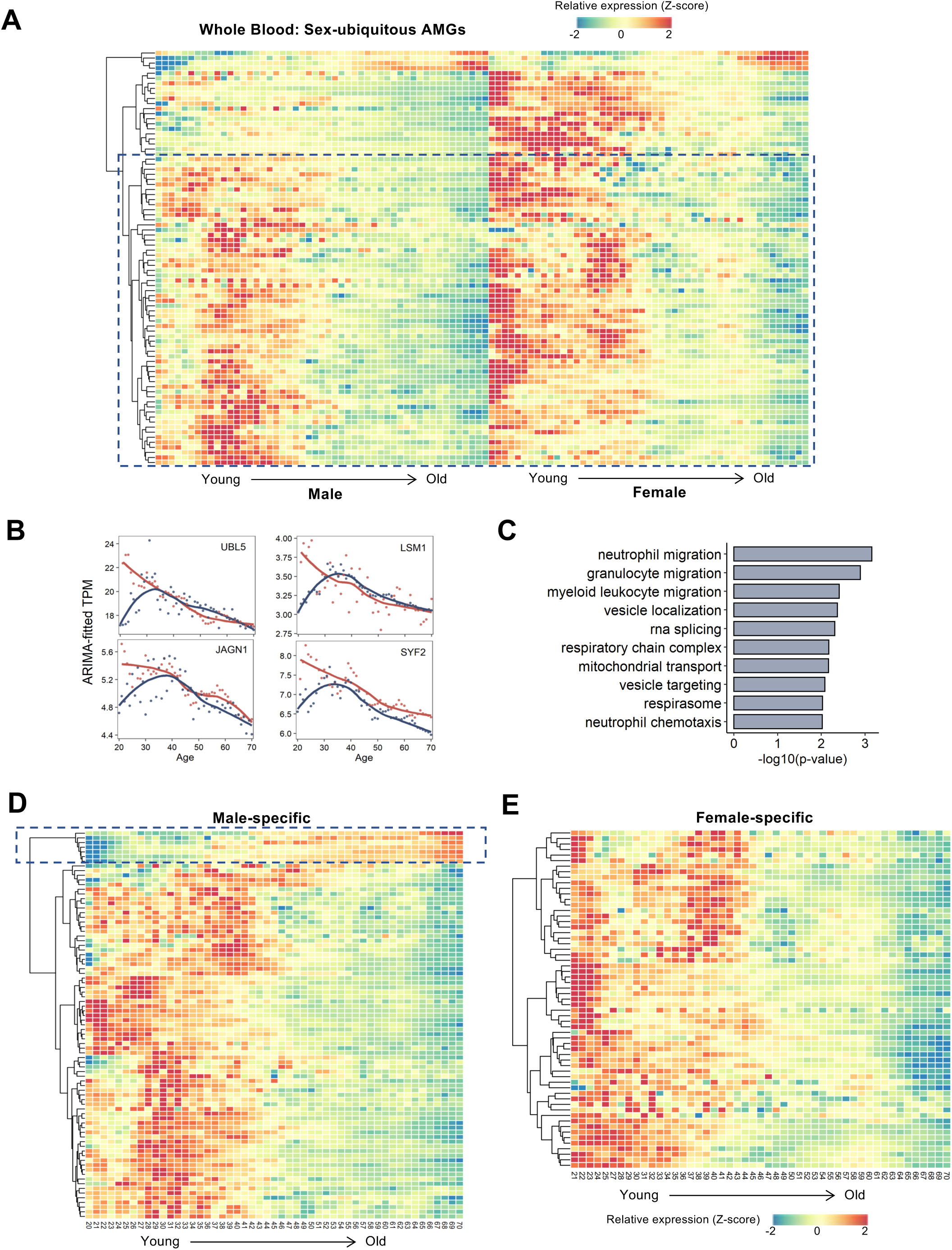
GE patterns of the AMGs in whole blood tissue.

