## Supplementary figures for "Pan-tissue Transcriptome Analysis Reveals Sex-dimorphic Human Aging"

**Fig. S1.**

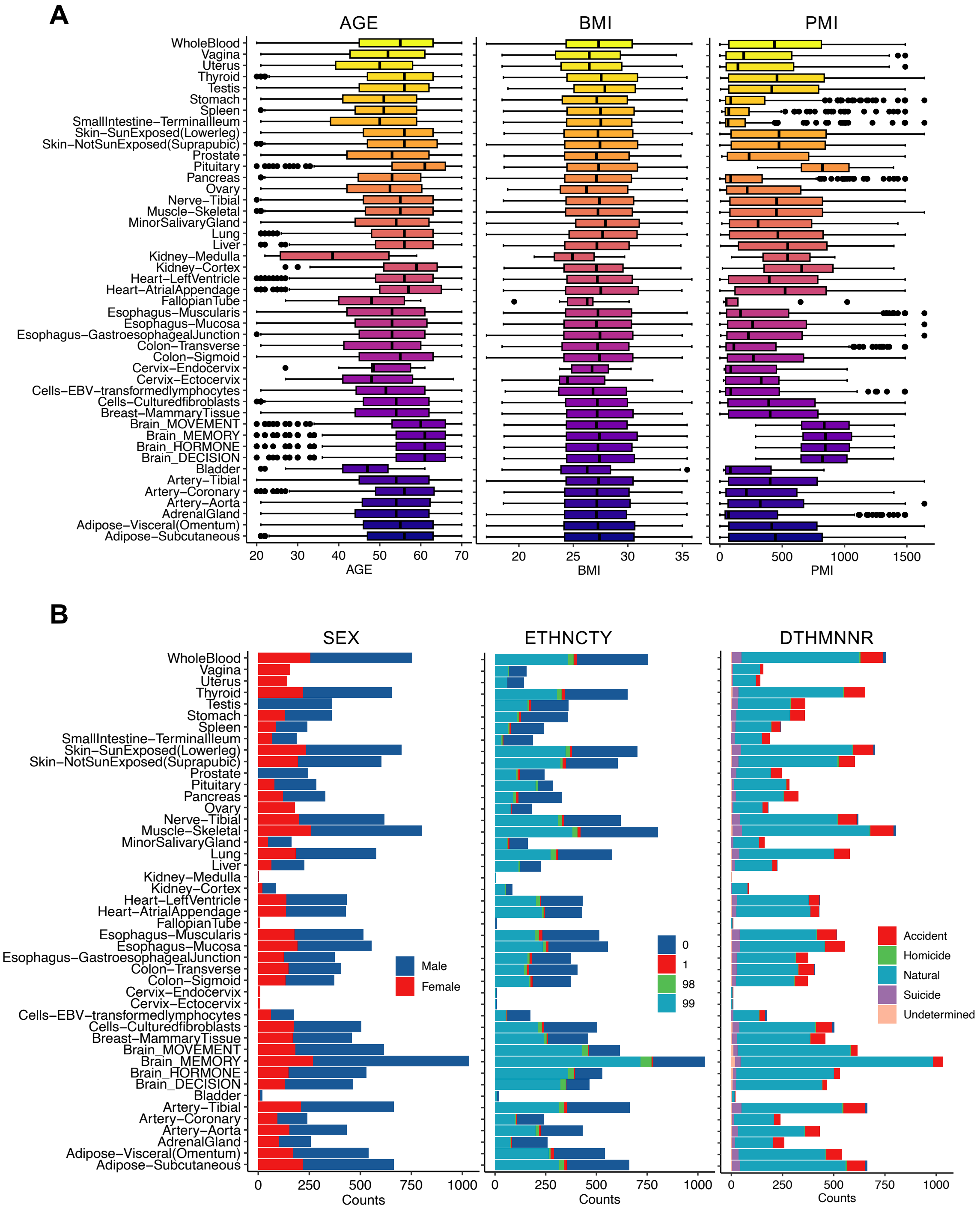


**Fig. S1 Sample phenotype in GTEx v8 cohort.**

**(A)** Age, BMI (Body Mass Index), and PMI (Post-Mortem Interval) distribution in GTEx v8 cohort.

**(B)** Sex, ethnicity, and manner of death (DTHMNNR) distribution in GTEx v8 cohort.

### Fig. S2.

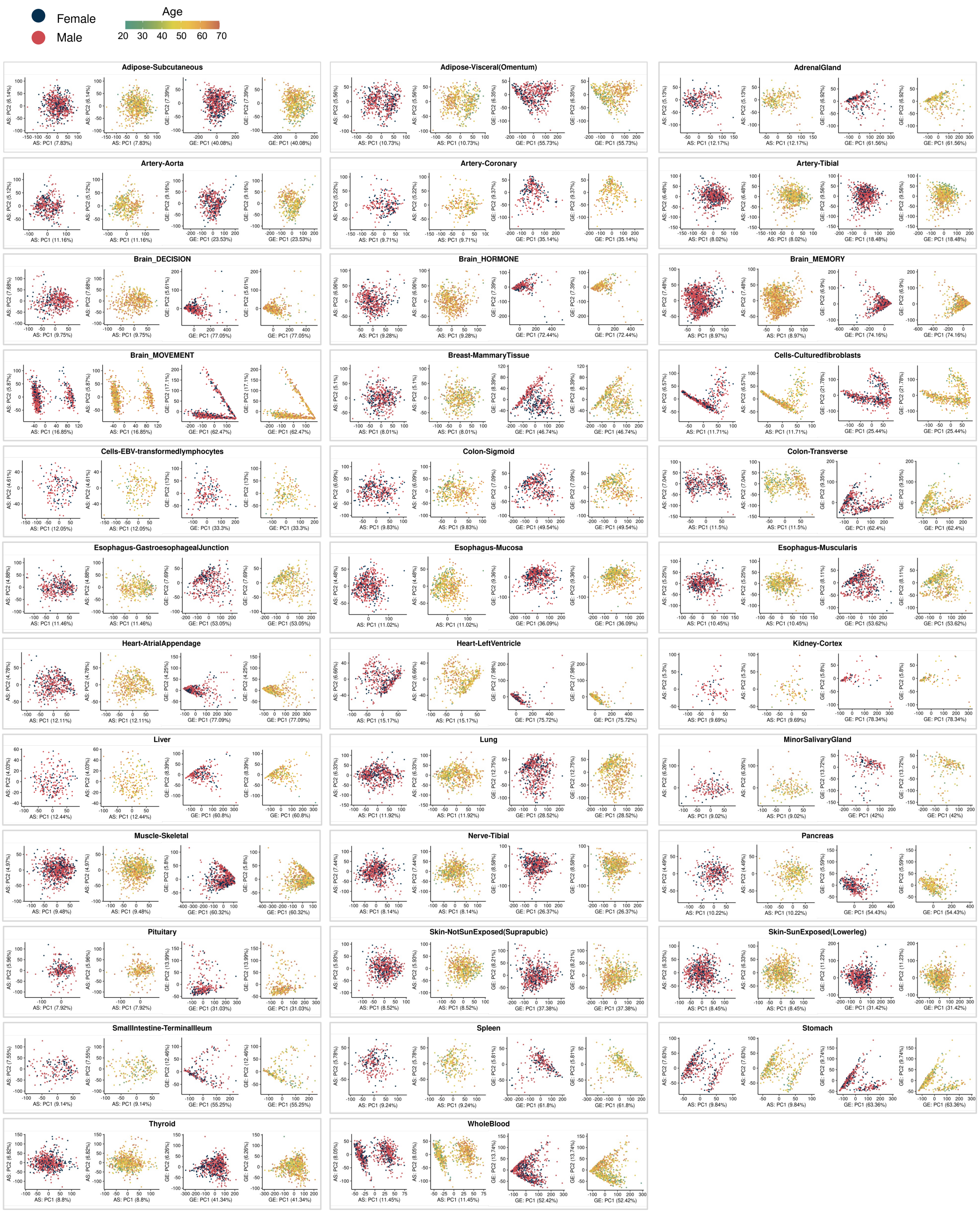


**Fig. S2 PCA on GE and AS profile.**
PCA is performed using GE and AS profiles, respectively. The X and Y-axis indicate scores of the top 2 PCs. Female and male samples are labeled with the red and blue dots, while samples with continuous ages are labeled in a gradient palette. The percentage of explained variation by each PC is labeled nearby each axis.

**Fig. S3.**

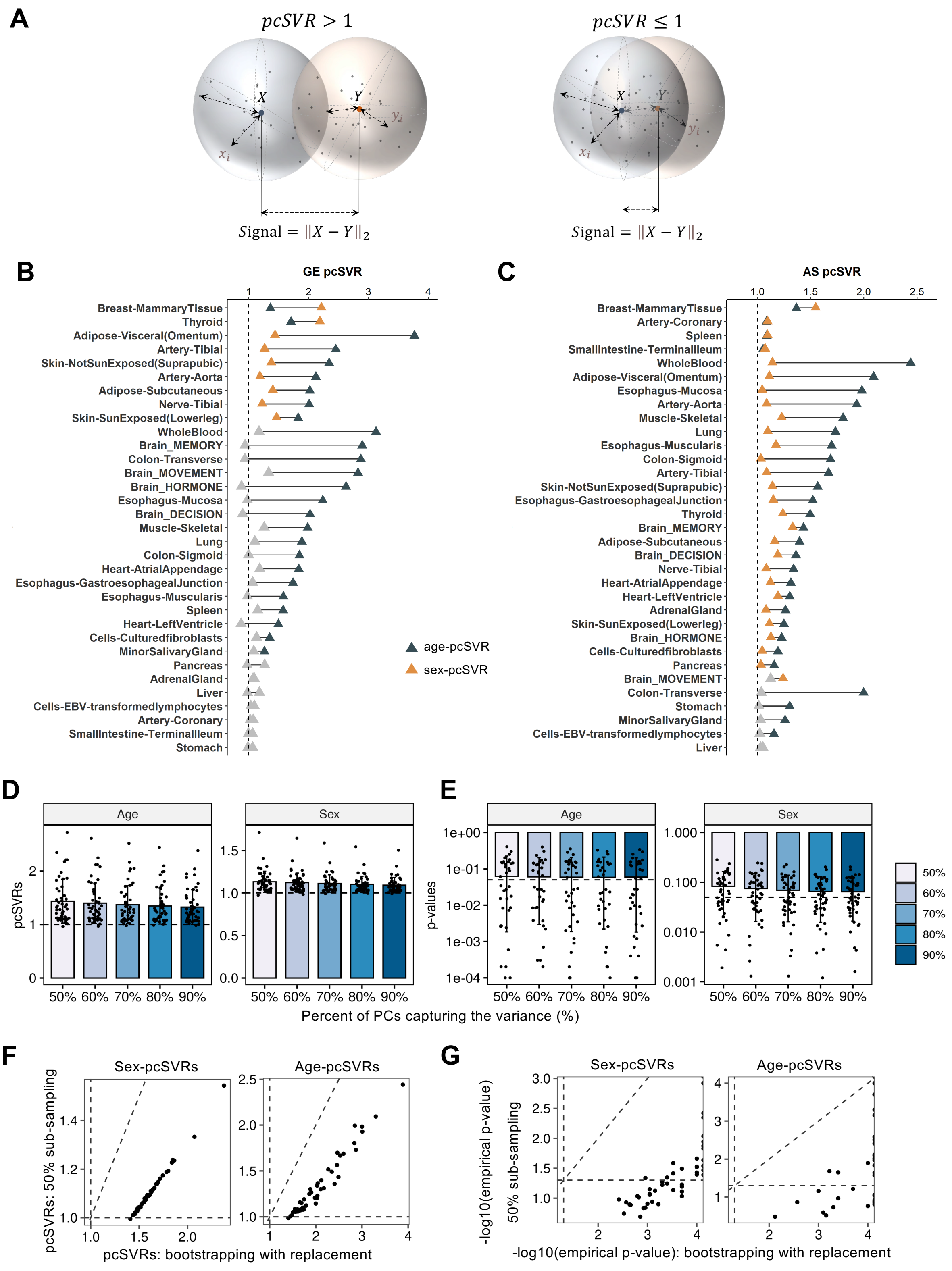


**Fig. S3 pcSVR calculation and robustness.**

**(A)** Schematic diagram of pcSVR calculation. pcSVR higher than 1 indicates that the differences between groups are higher than the noise within groups (left). pcSVR no more than 1 indicates that the differences between groups are smaller than the noise within groups (right). Two spheres represent two groups (females *vs.* males or young *vs.* old). Only three dimensions are shown.
**(B-C)** pcSVR between different sex or age groups calculated by GE (**B**) and AS (**C**) without genes and AS events on sex chromosomes. The yellow triangles show significant pcSVR calculated between females and males, while the black triangles show significant pcSVR calculated between young (i.e., 20-40 years old) and old (i.e., >60 years old) groups using permutation p-value <0.1. Insignificant values are labeled in grey.
**(D-E)** Bar plot of the age- and sex-pcSVR values (**D**) and corresponding empirical p-values (**E**) calculated using different numbers of PCs capturing 50% to 90% of the variance. The dashed lines indicate the thresholds for significant sex/age effects judged by the pcSVR analysis.
**(F)** Correlations of pcSVRs calculated by the bootstrap and sub-sampling approaches as shown in scatter plots of sex-pcSVR and age-pcSVR values. The X-axis shows the results from the bootstrap approach, while the Y-axis shows the results from permutation analysis using 50% sub-sampling. Each point indicates a tissue.
**(G)** Correlations of empirical p-values calculated by the bootstrap and sub-sampling approaches as shown in scatter plots of -log10 transformed empirical p-values. The label of X- and Y-axis is similar to panel **F**. The dots at the right margin indicate the empirical p-values equal to 0, with infinite -log10 transformed values.

**Fig. S4.**

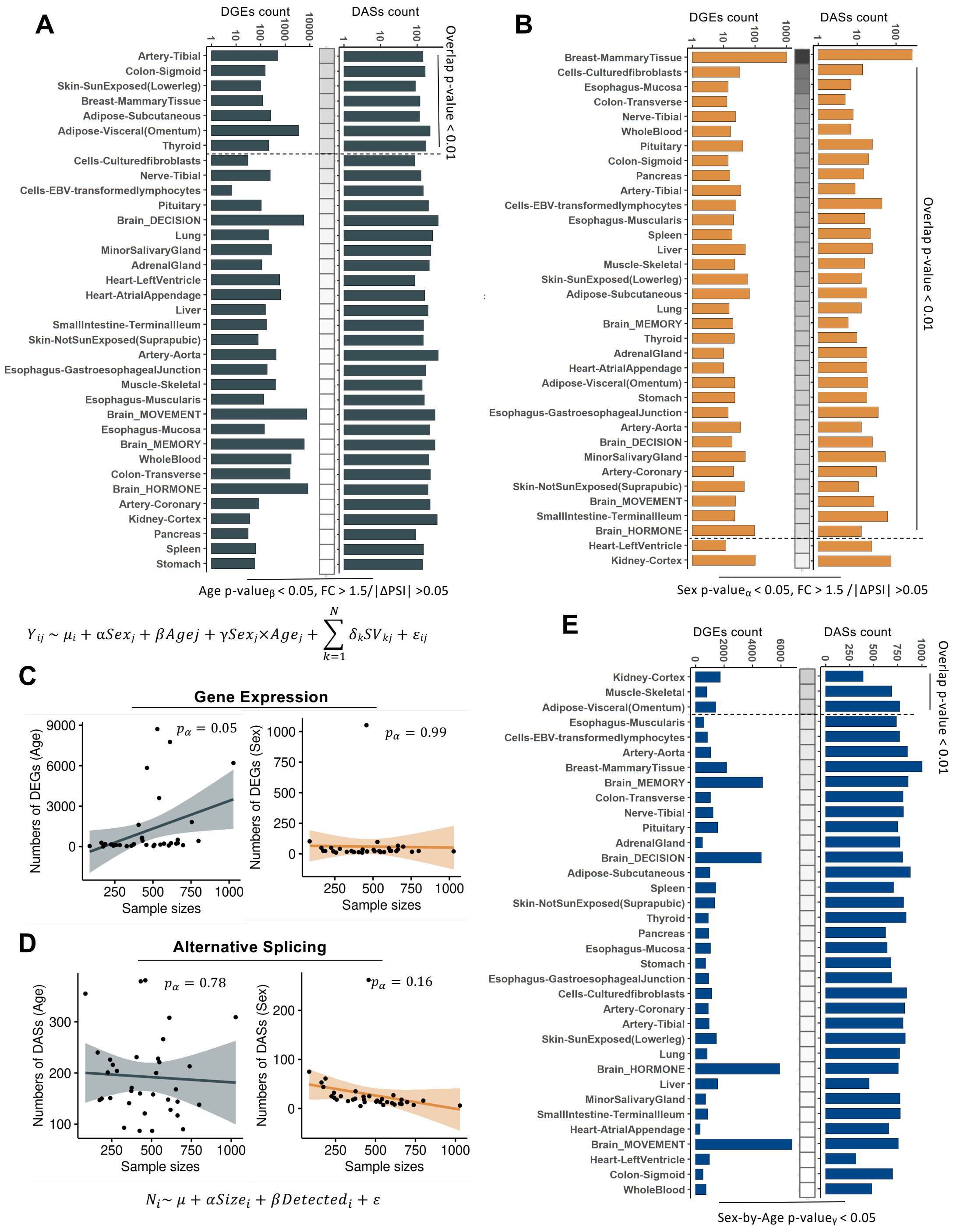


**Fig. S4 Differential GE and AS analysis to identify sex/age-differential genes and AS events.**

**(A-B)** The numbers of age-differential (**A**) and sex-differential(**B**) genes (left) and AS events (right). The color bar between the two bar plots represents the overlapped significance between sex/age-differential genes and spliced genes calculated using hypergeometric test. The dashed lines separate the tissues with significant or non-significant overlaps using the p-value <0.01. The sex-/age-differential genes and AS events are defined with coefficient p-value <0.05 and fold change >1.5 for GE or $\Delta$PSI >0.05 for AS.

**(C-D)** Correlation between the sample sizes and the numbers of age/sex-differential genes (**C**) or AS events (**D**). The X-axis indicates the total sample sizes. The Y-axis shows the numbers of age-differential genes/AS events (left) and sex-differential genes/AS events (right). The p-values of the coefficient of the sample sizes are calculated by linear regression as the formula on the bottom.

**(E)** The numbers of genes (left) and AS events (right) with significant age-by-sex interactions. The color bar between two bar plots represents the overlapped significance between genes and spliced genes calculated using hypergeometric test. The dashed lines separate the tissues with significant or non-significant overlaps.

**Fig. S5.**

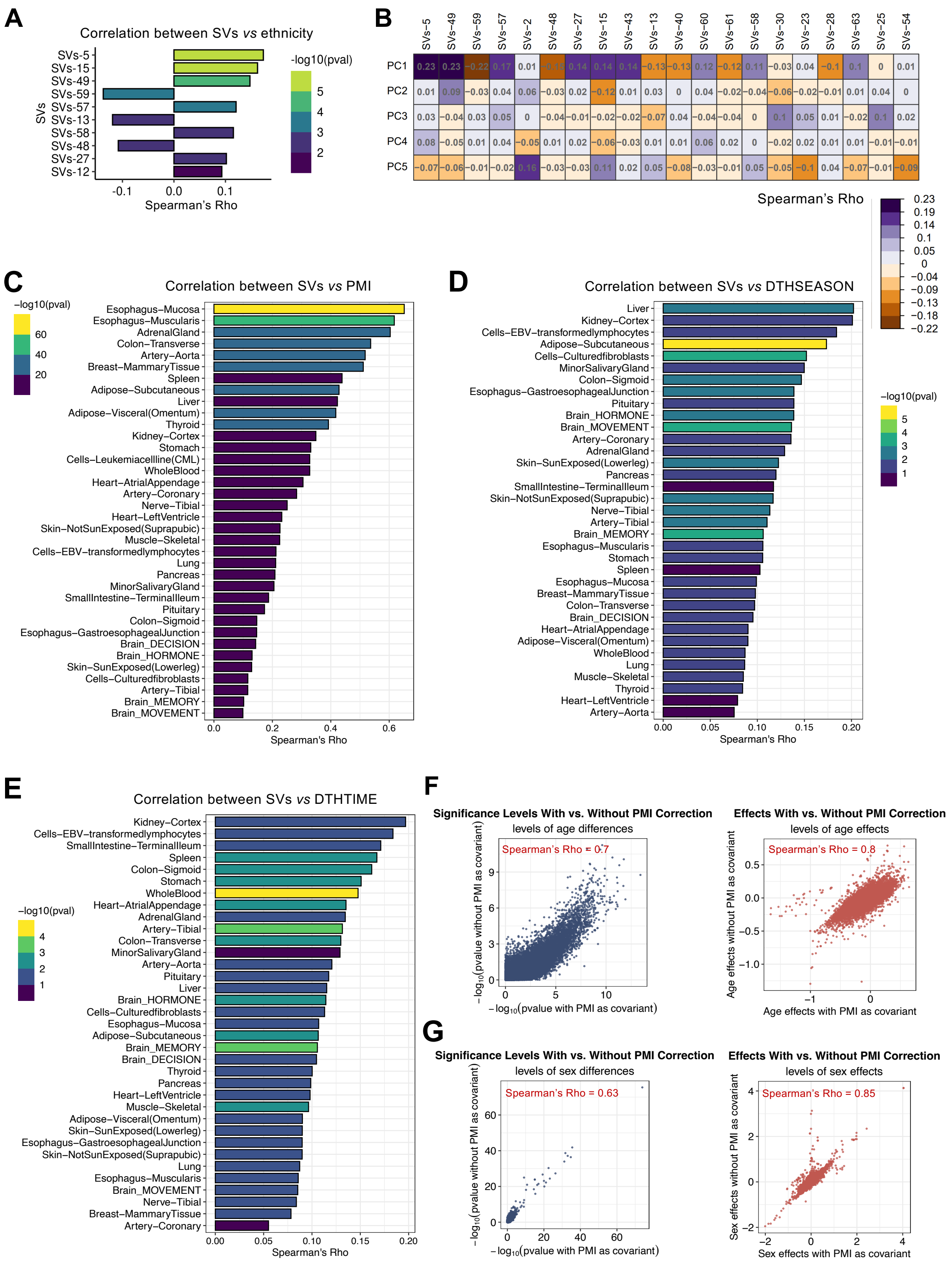


**Fig. S5 The evaluation of the confounding factors in the linear regression model.** (**A-B**) Correlation between the surrogate variables and human genetic background, including the donor ethnicity (**A**) and the principal components from whole genome sequencing (WGS) data (**B**).

(**C-E**) The highest correlation ρ between surrogate variables vs post-mortem interval (PMI) (**C**), season of death (**D**), and time of death (**E**). Each bar represents those correlations in each human tissue. The bars are color-coded according to a -log10(p-value) scale shown on the right.
(**F-G**) The results of differential gene expression analysis with vs without the inclusion of PMI correction as a known covariate. The scatter plots show the correlations of significance levels (p-values, left panel) and effect sizes (coefficients, right panel) of sex (**F**) and age (**G**). Whole-blood tissue is used as an example.

**Fig. S6.**

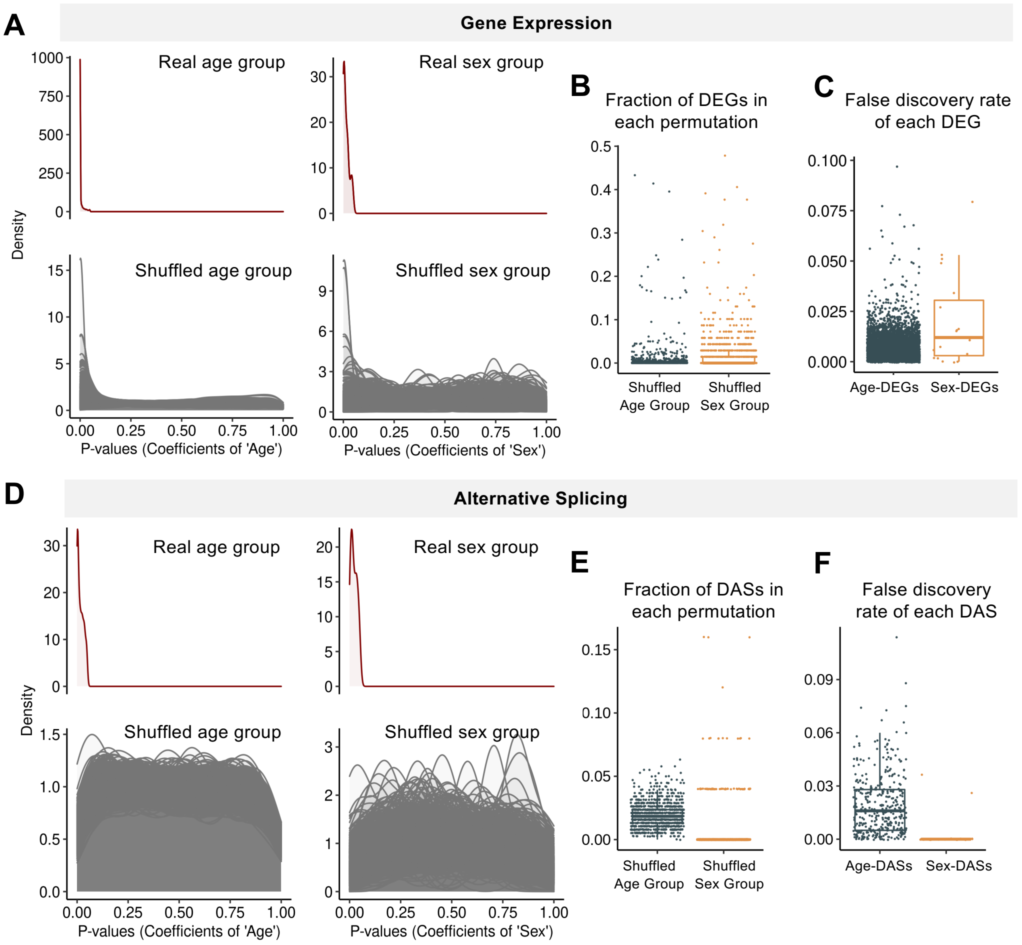


**Fig. S6 Permutation analysis of the linear regression model in differential analysis.**
(**A, D**) Distribution of the p-values across sex-/age-differential genes (**A**) or AS events (**D**) identified from original groups or shuffled groups in 1,000 permutated iterations.
(**B, E**) Fractions of original sex-/age-differential genes (**B**) or AS events (**E**) that are detected in each permutation. Each data point indicates the result from one shuffled permutation.
(**C, F**) False discovery rate (FDR) of each age-/sex-differential gene (**C**) and AS event (**F**) calculated across 1,000 permutations. Each data point indicates the result of one age-/sex-differential gene (**C**) and AS event (**F**).

**Fig. S7.**

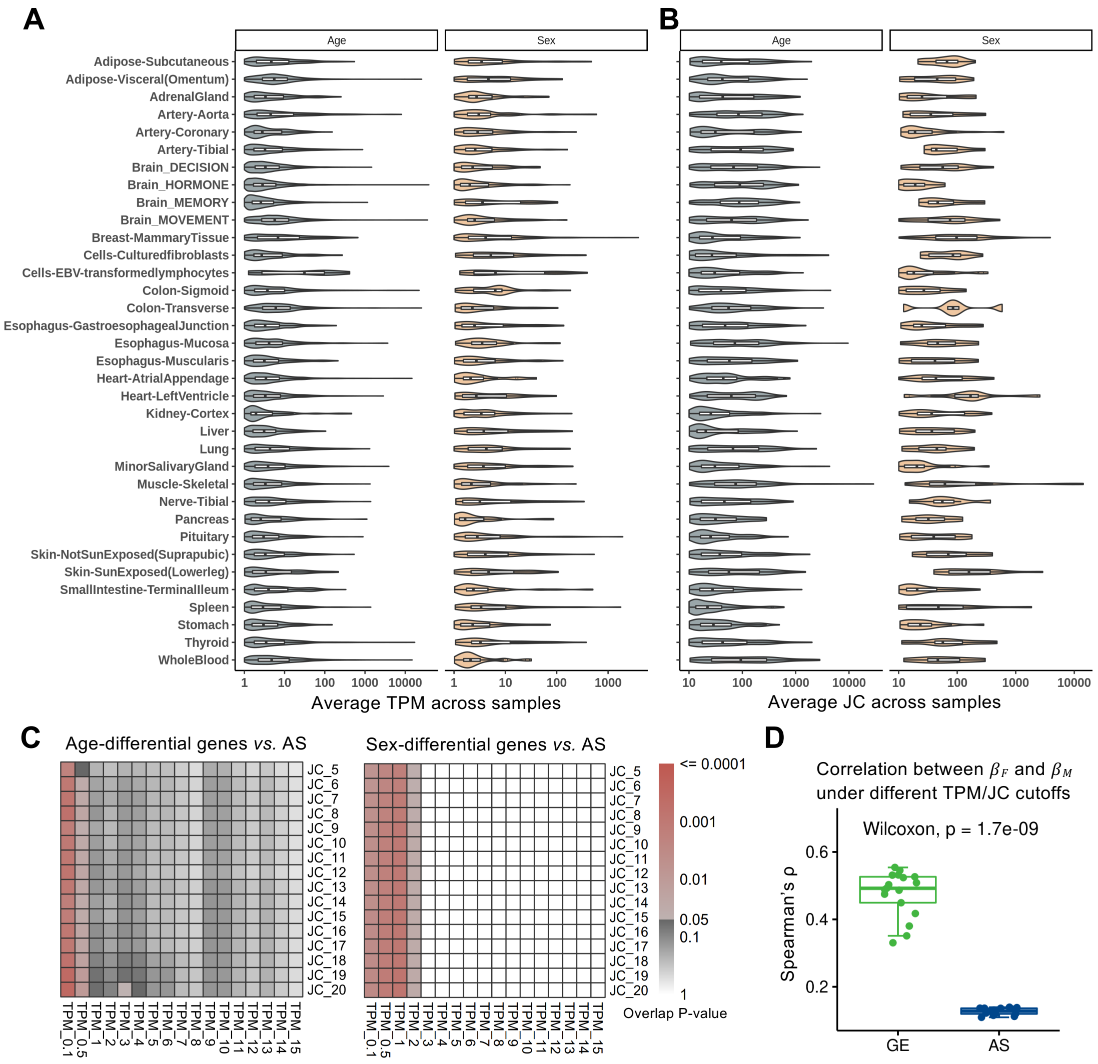


**Fig. S7 Robustness of GE and AS cutoffs.**

**(A-B)** The distribution of junction reads counts (**A**) and TPMs (**B**) of age-/sex-differential genes/AS events in multiple tissues. The X-axis indicates the average TPMs or JCs across the samples in certain tissue. The X-axis indicates the average TPMs or JCs across the samples in certain tissue.

**(C)** Overlap between the age/sex-differential genes and AS genes in decision-related brain region. The overlapped p-values calculated by hypergeometric test are shown with the color bar.

**(D)** Correlation of the age effects between females and males on GE (green) or AS (blue) using decision-related brain regions as example. Y-axis represents the correlations of the effect sizes of age ($\beta_F$, $\beta_M$) between two sexes across all genes/AS events (Spearman's correlation). Each jitter indicates the correlation under each single cutoff. P-values are estimated using Wilcoxon signed-rank test.

**Fig. S8.**

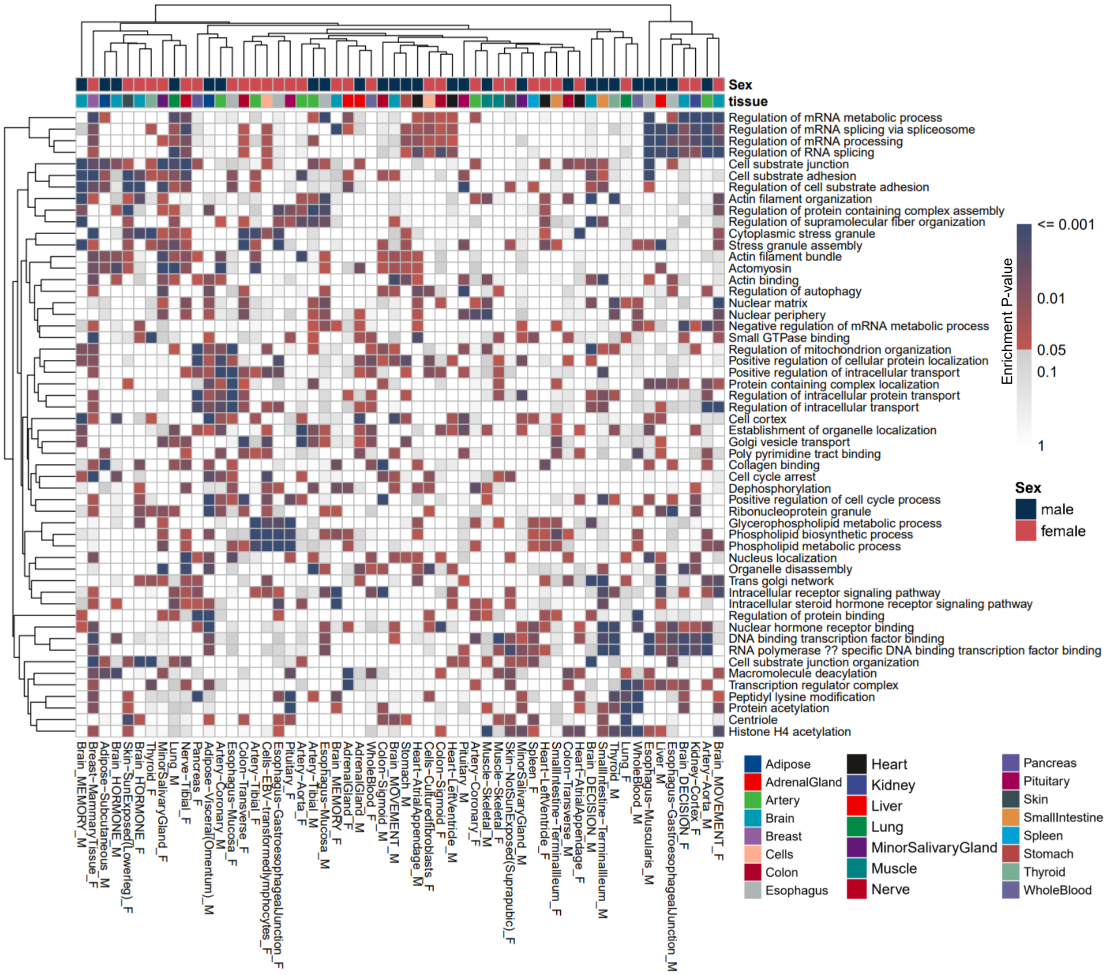


**Fig. S8 Functional enrichment of sBASEs across tissues.**
GO enrichment of the sBASEs in females or males across all tissues. Sex-specific GO terms significantly in at least 15 tissues were shown.

**Fig. S9.**

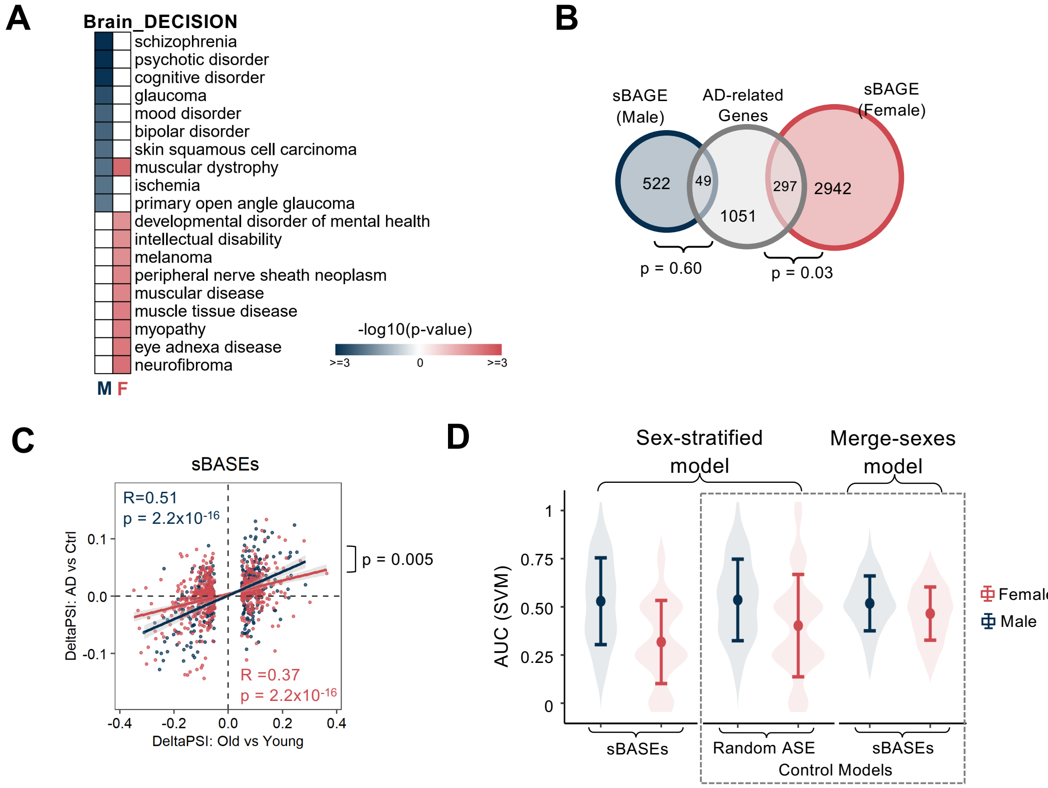


**Fig. S9 Sex-biased associations between sBASEs and diseases in decision-related brain region.**

**(A)** DO analysis of the sBASEs in females or males in decision-relation brain region. Sex-specific DO terms with top10 -log10 transformed enrichment p-values in each sex were shown.

**(B)** Venn diagram between sex-biased age-associated genes and AD-related genes. The p-values are calculated using hypergeometric test.

**(C)** Correlation between AD-associated and age-associated AS changes across the sBASEs in females and males from the ROSMAP dataset. The X-axis indicates the age-associated AS changes ($PSI_{Old}$-$PSI_{Young}$), while Y-axis indicates AD-associated AS changes ($PSI_{AD}$-$PSI_{Control}$).

(**D**) Performances of sex-stratified and merge-sexes models predicted by sBASEs and randomly selected AS events for 100 iterations using SVM classifier. The control models (i.e., the sex-stratified model trained by randomly selected AS events and the merge-sexes model trained by sBASEs in each sex) are highlighted with dashed lines. The mean ± sd (standard deviation) of AUC in 100 iterations is shown in the boxplot.

**Fig. S10.**

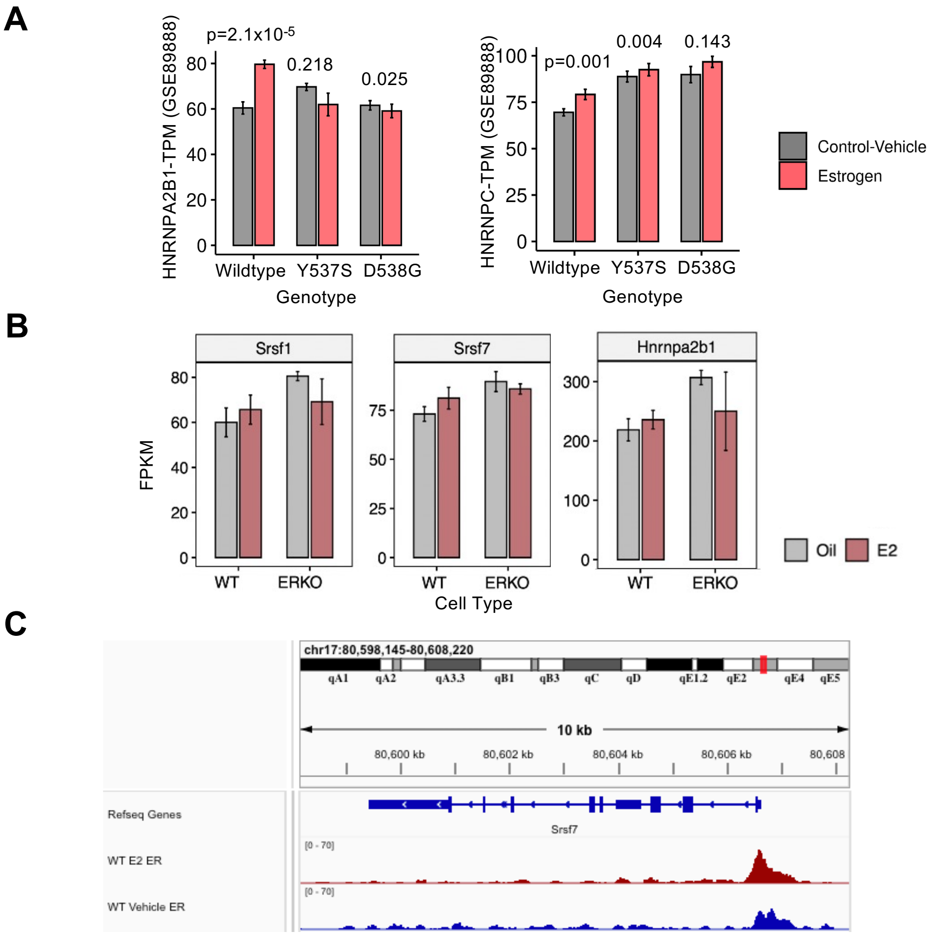


**Fig. S10 Gene expression regulation of sex-biased age-associated splicing factors by estrogen via ESR1 in multiple datasets.**

(**A**) Expression levels of HNRNPA2B1 and HNRNPC treated by 1nM estradiol (E2) vs. vehicle control (veh) in ESR1 wildtype, Y537S, and D538G mutant MCF-7 cell lines. Y-axis indicates the TPM values; the p-value are labeled on the top.

(**B**) Expression levels of Hnrnpa2b1, Hnrnpc, Srsf1 and Srsf7 treated by estradiol (E2) *vs.* vehicle control (Oil) of ERα wildtype and knockout ARCs in female mice. Y-axis indicates the FPKM values.

(**C**) ESR1 ChIP-seq reads coverages aruond Srsf1 and Srsf7 treated by estradiol (E2) *vs.* vehicle control (Oil) of ERα wild-type (WT, top panel) and knock-out (KO, bottom panel) in female mice. Y-axis of each track indicates the read counts at each position from ChIP-seq.

**Fig. S11.**

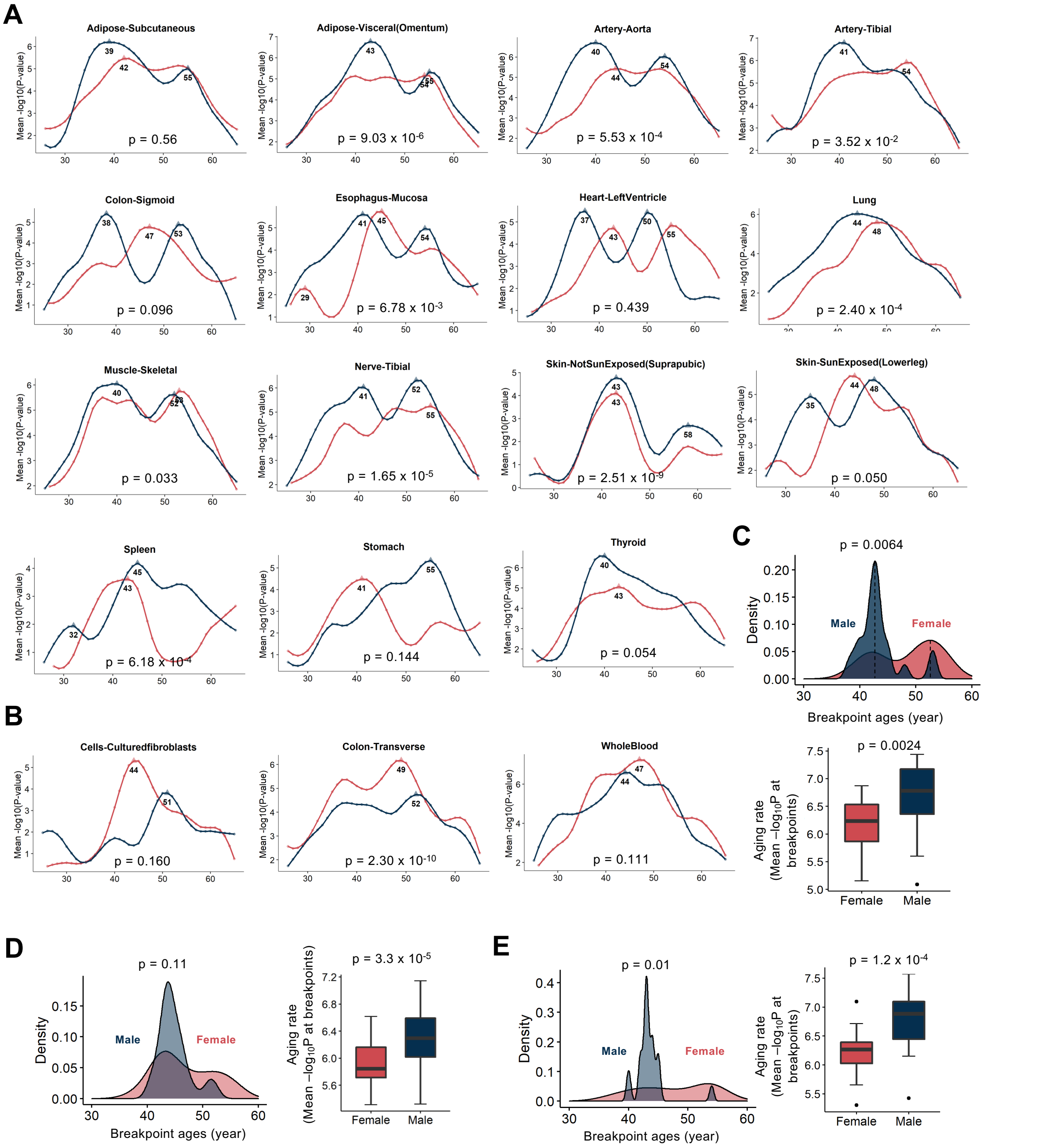


**Fig. S11 Breakpoint analysis across multiple tissues.**

**(A-B)** Tissues with faster aging rates in males (**A**) and females (**B**). The p-values are calculated using Wilcoxon signed-rank test and are labeled at the bottom.

**(C)** Characteristics of the major breakpoints calculated by all chronological genes in females and males. The left panel shows the distribution of the major breakpoints across multiple tissues. The right panel shows the aging rate at the major breakpoints. The p-values are both calculated using Wilcoxon signed-rank test.

**(D-E)** Characteristics of the major breakpoints calculated by chronological sBASEs (**D**) and all chronological AS events (**E**) in females and males. The left panel shows the distribution of the major breakpoints across multiple tissues. The right panel shows the aging rate at the major breakpoints. P-values are both calculated using Wilcoxon signed-rank test.

Fig. S12.

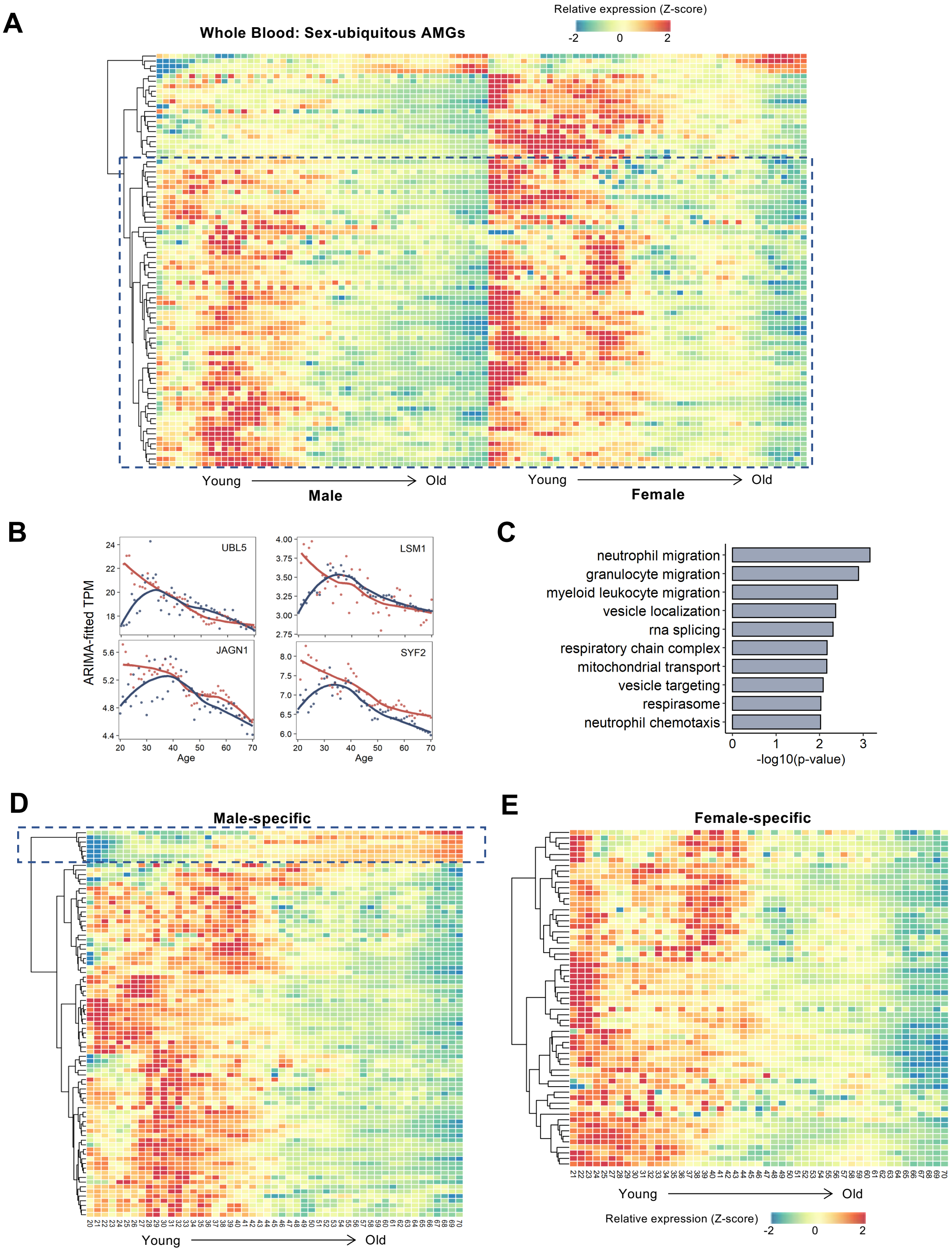


**Fig. S12 GE patterns of the AMGs in whole blood tissue.**
**(A)** GE patterns of AMGs common in both sexes (counts = 82) during aging. The values represent the z-score of fitted TPM from ARIMA models.

**(B)** Examples of AMGs common in both sexes with specific GE patterns. The dots indicate the ARIMA-fitted expression levels at each age point. The solid lines indicate the fitted values from LOESS regression for displaying the GE trends during aging.
**(C)** GO analysis of the AMGs common in both sexes with specific GE patterns during aging.
**(D-E)** GE patterns of male-biased (**D**, counts=83) and female-biased (**E**, counts=61) AMGs during aging. AMGs in dotted circles indicate the cluster of AMGs with increasing expression levels during aging in males.

**Captions for Table S1-S6**

**Table S1.**
Summary for sample sizes of each group in multiple tissues.

**Table S2.**
pcSVR values and corresponding permutation p-values between different age or sex groups as judged by GE and AS.

**Table S3.**
Lists of sex-stratified age-associated genes and AS events in 35 human tissues and brain regions.

**Table S4.**
P-values of GO analysis for sBASEs across multiple tissues.

**Table S5.**
AS regulatory networks between sBASEs and age-associated splicing factors in 4 functional brain regions.

**Table S6.**
Aging-modulated genes (AMGs) and enriched p-values across multiple tissues
